# Rho-dependent control of the Citron kinase, Sticky, drives midbody ring maturation

**DOI:** 10.1101/566463

**Authors:** Nour El-amine, Sabrya C. Carim, Denise Wernike, Gilles R.X. Hickson

## Abstract

Rho-dependent proteins control assembly of the cytokinetic contractile ring (CR), yet it remains unclear how those proteins guide ring closure and how they promote subsequent formation of a stable midbody ring (MR). Citron kinase is one important component required for MR formation but its mechanisms of action and relationship with Rho are controversial. Here, we conduct a structure-function analysis of the *Drosophila* Citron kinase, Sticky, in Schneider’s S2 cells. We define two separable and redundant RhoGEF/Pebble-dependent inputs into Sticky recruitment to the nascent MR and show that each input is subsequently required for retention at, and for the integrity of, the mature MR. The first input is via an actomyosin-independent interaction between Sticky and Anillin, a key scaffold also required for MR formation. The second input requires the Rho-binding domain of Sticky, whose boundaries we have defined. Collectively, these results show how MR biogenesis depends on the coordinated actions of Sticky, Anillin and Rho.

## Introduction

In animal cells, cytokinesis follows mitosis by forming a cleavage furrow at the equator of the dividing cell. Furrow ingression is driven at its base by a contractile ring (CR), a dynamic, membrane-anchored assembly of cytoskeletal proteins that constricts via disassembly to reduce the cell equator to a diameter of 1-2 µm (Schroeder, 1972; Carvalho *et al.*, 2009). At this point, the CR has gathered the bundled microtubules of the central-spindle, and it must transition into a stable structure called the midbody ring (MR). The MR and associated microtubule-based midbody form a dense structure (Mullins and Biesele, 1977), enriched in many different proteins and lipids (Skop *et al.*, 2004; Atilla-Gokcumen *et al.*, 2014). The CR-to-MR transition is a crucial step in cell division, essential for maintaining the integrity of the intercellular bridge and for setting the stage for the final abscission event that irrevocably separates the cells. The CR-to-MR transition occurs via a maturation process that includes a gradual thinning of the nascent MR and removal of cortical material through shedding and internalization (Mullins and Biesele, 1977; El Amine *et al.*, 2013; Renshaw *et al.*, 2014). The transition also involves changes in the actin and microtubule cytoskeleton (Hu *et al.*, 2012; Terry *et al.*, 2018). A complex machinery is involved in organizing the succession of events during cytokinesis, ensuring that they proceed with high fidelity (Fededa and Gerlich, 2012; Green *et al.*, 2012; D’Avino *et al.*, 2015; Glotzer, 2017). Activation of the small molecular weight GTPase RhoA is an early event essential for furrow initiation and CR formation (Bement *et al.*, 2005; Piekny *et al.*, 2005; Basant and Glotzer, 2018). The CR comprises filament-forming cytoskeletal proteins including actin, myosin II and septins, along with many other accessory proteins that ensure CR contractility, attachment to the plasma membrane, and interaction with central spindle microtubules (Green *et al.*, 2012; D’Avino *et al.*, 2015; Glotzer, 2017). Among these CR proteins is the Citron kinase, named Sticky in *Drosophila melanogaster*. Originally identified as a Rho-interacting protein (Madaule *et al.*, 1995; Di Cunto *et al.*, 1998; Madaule *et al.*, 1998), Citron kinase was initially suspected to play a role in myosin activation and cleavage furrow formation (Madaule *et al.*, 1998; Madaule *et al.*, 2000; Yamashiro *et al.*, 2003). However, it has since become clear that the major function and requirement for Citron kinase during cytokinesis is not at the CR *per se*, but rather in the formation of the MR that occurs after CR closure (D’Avino *et al.*, 2004; Echard *et al.*, 2004; Naim *et al.*, 2004; Dean and Spudich, 2006). Although there is evidence of redundant contributions of Citron kinase/Sticky during furrowing (Madaule *et al.*, 1998; Gai *et al.*, 2011; El Amine *et al.*, 2013), as well as suggestions of a role during abscission (Naim *et al.*, 2004; Gai *et al.*, 2011; Sgro *et al.*, 2016), loss-of-function studies have clearly shown that Citron kinase/Sticky is required for the stabilization of the MR in a variety of cell types (Echard *et al.*, 2004; Gruneberg *et al.*, 2006; Neumann *et al.*, 2010; Gai *et al.*, 2011; Bassi *et al.*, 2013; El Amine *et al.*, 2013; Watanabe *et al.*, 2013; McKenzie *et al.*, 2016). However, precisely how Sticky is regulated and how it acts to stabilize the MR remains unclear. Although Citron kinase is primarily a cortical protein, numerous studies indicate that it can interact with microtubule-associated proteins of the midbody, such as KIF14/Nebbish, kinesin-6/MKLP1/Pavarotti and PRC1, and that these interactions are important for intracellular bridge stability (Gruneberg *et al.*, 2006; Bassi *et al.*, 2013; Watanabe *et al.*, 2013; McKenzie *et al.*, 2016).

An important and longstanding question concerns the nature of the relationship between Citron kinase/Sticky and the Rho GTPase. The original assumption that Sticky acts as a canonical downstream effector of Rho has recently been challenged by the suggestion that mammalian Citron kinase acts upstream of RhoA to modulate its activity (Gai *et al.*, 2011; Dema *et al.*, 2018). Additionally, there is evidence of *Drosophila* Sticky interacting with Rho via its C-terminal Citron-Nik1 homology (CNH) domain, and not via its putative Rho-binding domain (Shandala *et al.*, 2004; Bassi *et al.*, 2011; Gai *et al.*, 2011). Thus, the relationship between Citron kinase and Rho remains controversial and unclear.

Here, we show in *Drosophila* S2 cells that the RhoGEF Pebble controls the localization of Sticky to the cleavage furrow through two partially redundant and separable inputs, acting on its central coiled-coil region. One input depends on the highly conserved Sticky RBH region and likely involves direct binding of Rho1-GTP, while the second input requires interaction with the Rho1-dependent scaffold protein, Anillin, also known as Scraps in *Drosophila*. Disrupting either the Rho1/Sticky or the Anillin/Sticky interaction led to failures in MR formation, indicating that both inputs are required for Sticky function. Conversely, little to no evidence was found to support a role for the CNH domain in Sticky localization or function during cytokinesis. Collectively, the data lead to the proposal that Rho1 acts on Sticky both directly, and indirectly via Anillin, to recruit Sticky to the CR. In turn, Sticky acts to retain Anillin in the subsequent MR as it forms around the midbody, to which Sticky is also connected. Thus, Rho1 controls Sticky through multiple inputs to coordinate MR formation.

## Results

### Sticky localization requires Rho1 signaling

To test the requirements for Rho1 signaling in the localization of Sticky, we first depleted the RhoGEF Pebble (the ortholog of mammalian ECT2) in cells expressing Sticky-GFP. We note that many Sticky-derived constructs exhibit cytoplasmic punctate structures, presumed to be aggregates, as described for mammalian Citron kinase (Eda *et al.*, 2001; Watanabe *et al.*, 2013). Nevertheless, non-aggregated Sticky-GFP is still robustly recruited to the cell cortex during cytokinesis and, as previously shown, is able to functionally rescue for loss of endogenous Sticky (El Amine *et al.*, 2013). Live-cell spinning disc confocal microscopy showed that cells incubated with control *lacI* dsRNA (Figure 1B, Supplemental Movie 1) or a dsRNA targeting the Sticky 3’UTR (data not shown, but see Supplemental Figure S1 for efficacy of different Sticky dsRNAs used in this study), robustly recruited Sticky-GFP to the CR and MR, and exhibited some shedding from the nascent MR, similar to that previously described for Anillin (El Amine *et al.*, 2013). However, in cells incubated with Pebble (*pbl*) dsRNA, Sticky-GFP localization to the cell cortex was abolished during anaphase/telophase, indicating that Pebble controls Sticky recruitment (Figure 1C). Sticky contains a highly conserved, 28-amino acid, Rho-binding homology (RBH) region within its central coiled-coil region (residues 1235-1263) that is predicted to bind to Rho-GTP (Bassi *et al.*, 2011). Deletion of the RBH region (Sticky-ΔRBH-GFP, Figure 1D) or introduction of a point mutation (L1246N) predicted to abrogate any Rho-GTP binding (Shimizu *et al.*, 2003; Dvorsky *et al.*, 2004; Watanabe *et al.*, 2013; Jungas *et al.*, 2016) did not prevent cortical recruitment of Sticky (Figure 1E). However, live-imaging of deplete-and-rescue experiments indicated that these perturbations of the RBH greatly reduced the ability of Sticky to support cytokinesis (Figure 1H, N=68, \*\**p*=0.004 and N=87, \**p*=0.028 for Sticky-ΔRBH and Sticky-L1246N respectively). Thus, the RhoGEF/Pebble is required for Sticky localization and the RBH region is required for Sticky function, but is dispensable for cortical localization.

**Figure 1.**
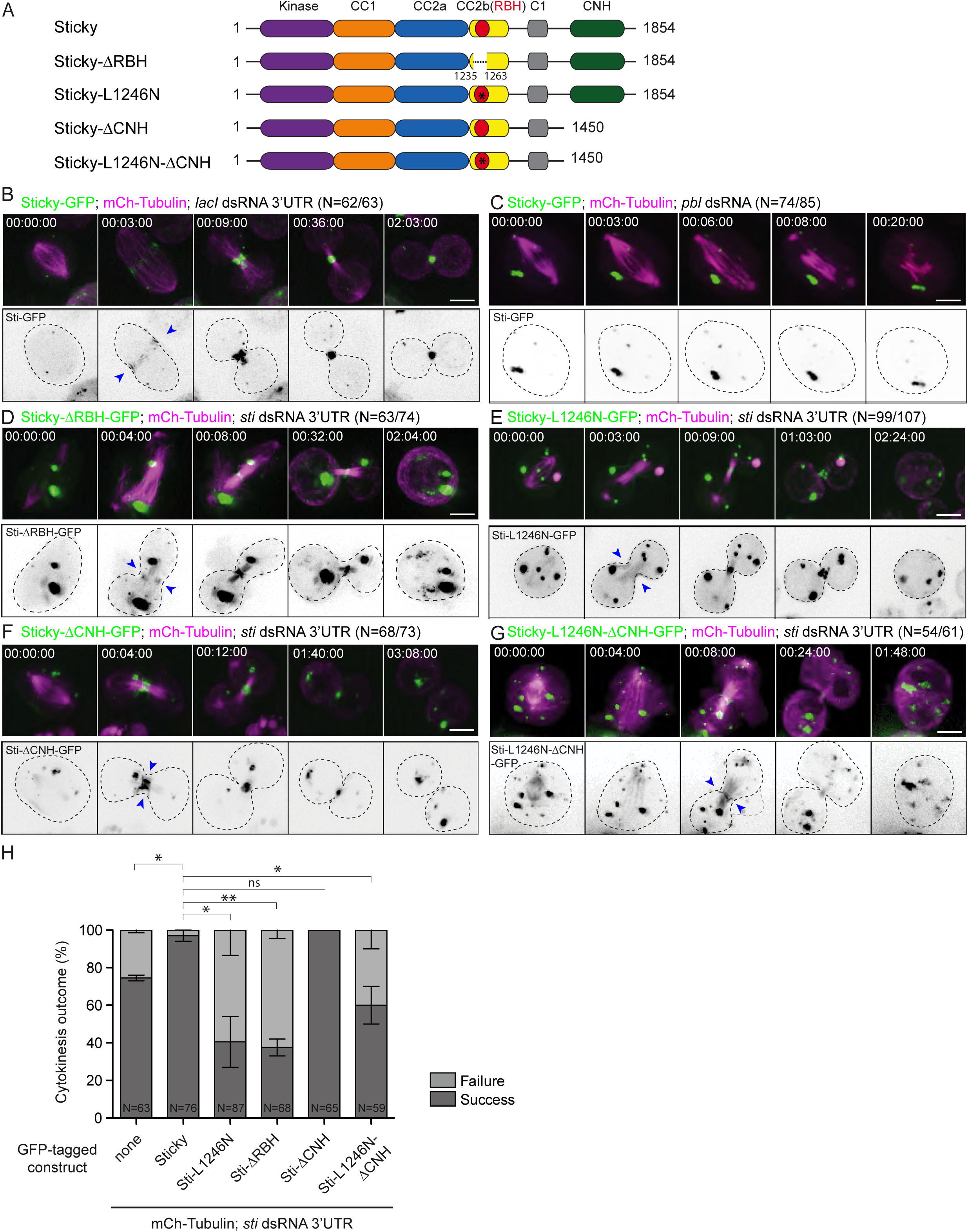
Sticky localization requires Rho1 signaling. A A cartoon representation of Sticky showing different domains, truncated constructs and the position of the L1246N mutation (asterisk). **B-F** Representative, high-resolution time-lapse sequences of cells stably expressing mCherry-Tubulin (magenta) and inducibly expressing (in green in the merged upper panels, or inverted grayscale in the lower panels) Sticky-GFP (A-B), Sticky-ΔRBH-GFP (C), Sticky-L1246N-GFP (D), Sticky-ΔCNH-GFP (E) or Sticky-L1246N-ΔCNH-GFP (F) following 3-day incubation with the specified dsRNAs. N represents the number of cells displaying similar patterns of GFP localization at the division plane at t=00:08:00-00:10:00/total number of cells scored. Data are from 2 independent experiments. Blue arrowheads highlight the ingressing cleavage furrow and dashed lines represent the outline of the cell in the grayscale images. **H** Quantification of the outcome (success or failure) of division attempts scored from low-resolution imaging of cells stably expressing mCherry-Tubulin and induced to express the specified GFP-tagged rescue constructs, following 3-day incubation with *sti* 3’UTR dsRNA. Data are from 2 independent experiments. Error bars represent standard deviation between experiments resulting from an unpaired t-test with \**p* < 0.05; \*\**p* < 0.01 for significance and ns = non-significance. Times are in h:min:sand scale bars are 5 µm.

The C-terminal CNH domain has also been reported to interact with Rho (Shandala *et al.*, 2004; Bassi *et al.*, 2011). However, a construct lacking this domain (Sticky-ΔCNH-GFP; residues 1-1450) was indistinguishable from full-length Sticky-GFP in terms of localization to the CR and MR (Figure 1F) and in its ability to support successful cytokinesis upon depletion of endogenous Sticky via *sti* dsRNA 3’UTR (100% success; Figure 1H, N=65, *p*=0.211). A construct combining deletion of the CNH domain with mutation of the RBH (Sticky-L1246N-ΔCNH-GFP) behaved similarly to Sticky-L1246N-GFP (60% vs. 41% success; Figure 1G-H, N=59, \**p*=0.036). We also tested the ability of the ΔCNH constructs to rescue the more penetrant failures induced by *sti* dsRNA3 that targets the CNH region (56% vs. 70% success, Supplemental Figure S2F, N=52 *p*=0.053). In this case, partial rescue by Sticky-ΔCNH-GFP was observed, in a manner that was abolished by the L1246N mutation (70% vs. 52% success; Supplemental Figure S2D, F, N=60, *p*=0.052). These results suggest that the CNH domain of Sticky is not essential for cytokinesis in S2 cells. However, given that the rescue observed upon induction of Sticky-ΔCNH-GFP was incomplete (70% success vs. 56% in cells only expressing mCh-Tubulin, Supplemental Figure S2F), we cannot rule out the possibility that it plays a contributory role herein, or even an essential role in other cell types.

### Anillin contributes to the Rho-dependent cortical recruitment of Sticky

We previously showed that Sticky and Anillin cooperate during formation of the stable MR, and in particular that Sticky acts to retain Anillin at the nascent MR (El Amine *et al.*, 2013). We therefore wished to test whether Anillin, itself a Rho-dependent protein (Hickson and O’Farrell, 2008b; Piekny and Glotzer, 2008; Sun *et al.*, 2015), contributes to the cortical localization of Sticky. Depletion of Anillin using different, efficacious dsRNAs (Supplemental Figure. S4) that block complete closure of the CR, did not prevent the cortical recruitment of Sticky-GFP, indicating that Anillin is dispensable for Sticky localization (Figure 2B, arrowheads at 00:03:00 and 00:09:00). However, Anillin depletion did prevent the cortical recruitment of Sticky-L1246N-GFP (Figure 2C, Supplemental Movie 2) and Sticky-ΔRBH-GFP (Figure 2D), albeit Sticky-ΔCNH-GFP was still recruited following Anillin depletion (Figure 2E, arrowheads at 00:05:00 and 00:15:00). These results indicate that Sticky recruitment to the CR involves at least 2 inputs: one dependent on Anillin; the other dependent on Sticky’s RBH, and with no major contribution from the CNH domain.

**Figure 2.**
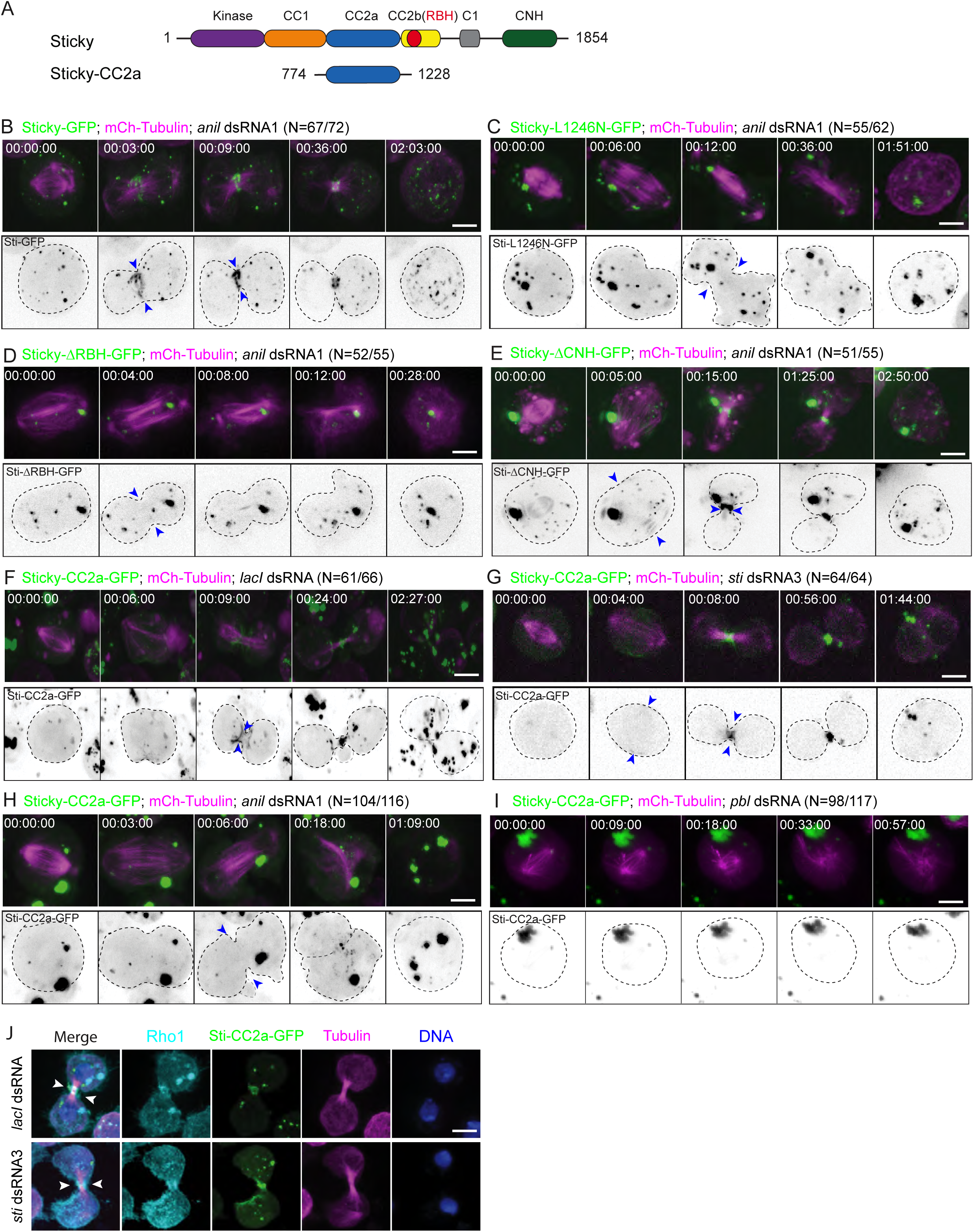
Anillin contributes to the Rho1-dependent cortical recruitment of Sticky. A A cartoon representation of Sticky showing different domains and Sticky-CC2a. **B-I** Representative, high-resolution time-lapse sequences of cells stably expressing mCherry-Tubulin (magenta) and inducibly expressing (in green in the merged upper panels, or inverted grayscale in the lower panels) Sticky-GFP (B), Sticky-L1246N-GFP (C), Sticky-ΔRBH-GFP (D), Sticky-ΔCNH-GFP (E) or Sticky-CC2a-GFP (F-I) following 3-day incubation with the specified dsRNAs. N represents the number of cells displaying similar patterns of GFP localization at the division plane at t=00:08:00-00:10:00/total number of cells scored. Data are from 2 independent experiments. Blue arrowheads highlight the ingressing cleavage furrow and dashed lines represent the outline of the cell in the grayscale images. **J** Immunofluorescence micrographs of S2 cells induced to express Sticky-CC2a-GFP, treated for 3 days with the indicated dsRNAs, fixed and stained with anti-Rho1, anti-Tubulin antibodies and Hoechst DNA stain. White arrowheads highlight the ingressed cleavage furrow. Times are h:min:s; scale bars are 5 µm.

Bassi *et al.* (2011) used bioinformatic approaches to subdivide the central coiled-coil region of Sticky into three sub-regions: CC1, CC2a and CC2b, and showed that CC2a (residues 774-1227), was sufficient for cortical localization. In agreement, we found that Sticky-CC2a-GFP (residues 774-1228) was robustly recruited to the CR and nascent MR (Figure 2F). The localization pattern of Sticky-CC2a-GFP remained unchanged upon RNAi-mediated depletion of endogenous Sticky (*sti* dsRNA3; Figure 2G), but it was abolished by depletion of Anillin (Figure 2H). RNAi of Pebble also prevented the recruitment of Sticky-CC2a-GFP, confirming that CC2a recruitment is RhoGEF-dependent (*pbl* dsRNA; Figure 2I). Immunofluorescence analysis of fixed S2 cells depleted of endogenous Sticky (*sti* dsRNA3) revealed co-localization of Sticky-CC2a-GFP and endogenous Rho1 at the nascent MR stage, providing further support that the recruitment of Sticky-CC2a is Rho1-dependent (Fig. 2J).

### The coiled-coil domain of Sticky contains two Pebble-dependent inputs

The results above suggested that Anillin contributes to Sticky localization via the CC2a region, while the RBH domain of Sticky, which resides within the CC2b region, also plays a role. To further test this notion, we closely examined the behavior of a construct containing both CC2a and CC2b (residues 774-1370) (Figure 3A), using live-cell imaging of mitotic cells. In cells treated with a control *lacI* dsRNA, Sticky-CC2a-CC2b-GFP was robustly recruited to the CR and nascent MR (Figure 3B). This recruitment was also seen upon depletion of endogenous Sticky, either using *sti* dsRNA 3’UTR (data not shown), or dsRNA3 that targets the mRNA sequence encoding part of the CNH (Supplemental Figure S1A), although the construct was unable to rescue cytokinesis failures (Figure 3C). As with full-length Sticky-GFP, Pebble depletion abolished the equatorial recruitment of Sticky-CC2a-CC2b-GFP (Figure 3D), whereas Anillin depletion did not (Figure 3E). Introduction of the L1246N mutation within the RBH domain did not prevent the cortical recruitment of Sticky-CC2a-CC2b-GFP (Figure 3F), unless Anillin was simultaneously depleted (Figure 3G). These observations confirm that the coiled-coil region of Sticky responds to two Pebble-dependent inputs: one via the CC2a region involving Anillin, and one via the RBH of Sticky within the CC2b region presumably involving Rho-binding. Furthermore, since both Anillin depletion and mutation of the RBH are required to abrogate localization, we conclude that these inputs act in a partially redundant fashion to recruit Sticky to the cortex.

**Figure 3.**
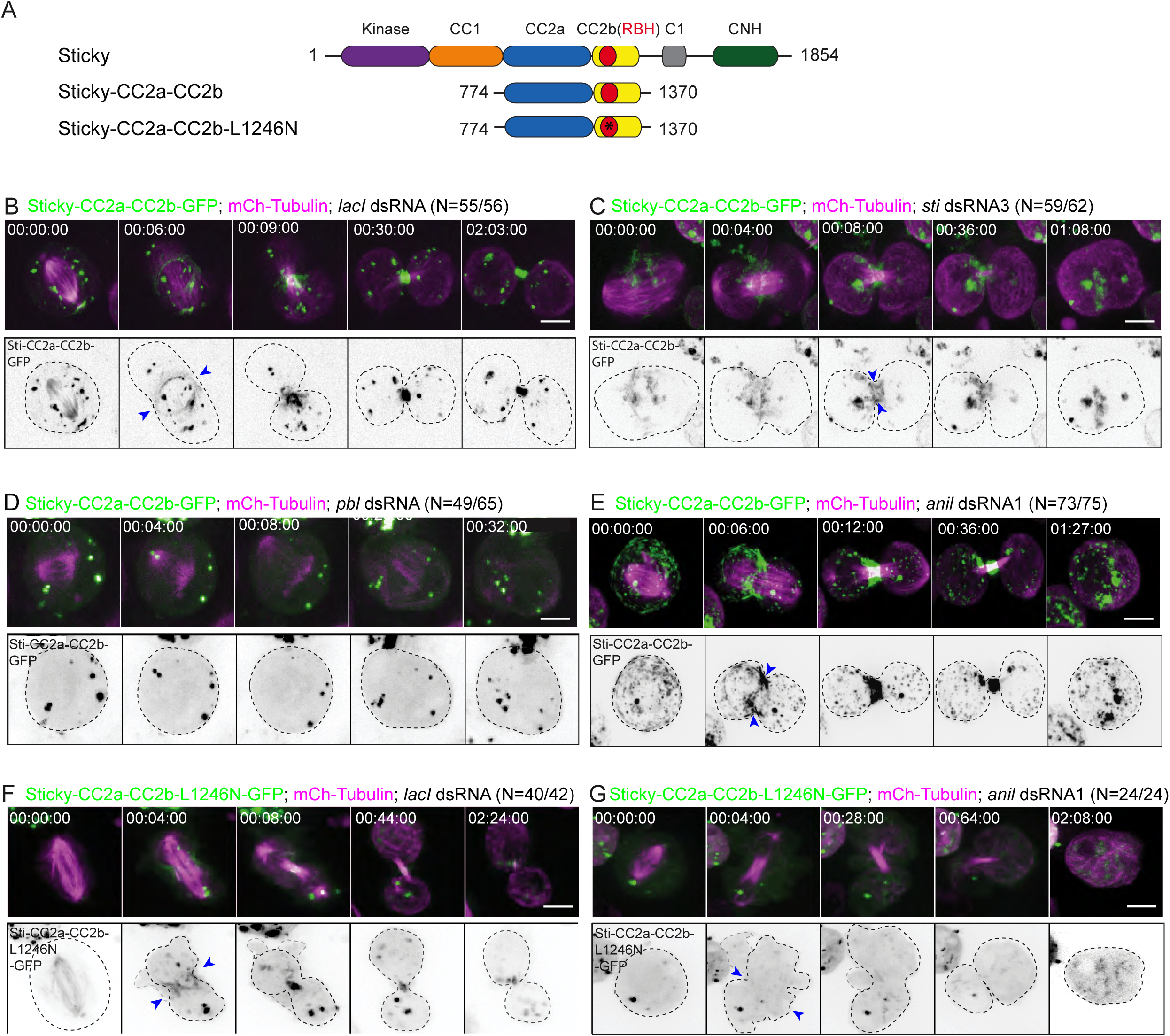
The coiled-coil domain of Sticky contains two Pebble-dependent inputs. **A** A cartoon representation of Sticky showing different domains and Sticky-CC2a-CC2b with and without the L1246N mutation (asterisk). **B-G** Representative, high-resolution time-lapse sequences of cells stably expressing mCherry-Tubulin (magenta) and inducibly expressing (in green in the merged upper panels, or inverted grayscale in the lower panels) Sticky-CC2a-CC2b-GFP (B-E) or Sticky-CC2a-CC2b-L1246N-GFP (F-G) following 3-day incubation with the specified dsRNAs. N represents the number of cells displaying similar patterns of GFP localization at the division plane at t=00:08:00-00:10:00/total number of cells scored. Data are from 2 independent experiments. Blue arrowheads highlight the ingressing cleavage furrow and dashed lines indicate the outline of the cell in the grayscale images. Times are h:min:s; scale bars are 5 µm.

### Defining the minimal Anillin-dependent input to Sticky localization

We sought to define the minimal requirements for each of these two inputs (CC2a and CC2b) and determine whether they could both be physically separated from one another. A series of constructs was generated in which the CC2a region (residues 774-1228) was progressively truncated by 50 residue increments from either the N- or C-terminus (Figure 4A-B). These fragments were fused to GFP, expressed in S2 cells, and their ability to localize to the furrow cortex was assessed by live-imaging time-lapse microscopy. Truncation of the N-terminus did not prevent cortical recruitment until residues 974-1024 were deleted (Figure 4A); the construct Sticky^1024-1228^-GFP and smaller constructs remained cytoplasmic during furrow ingression. Truncation of the C-terminus did not prevent cortical recruitment until residues 1078-1128 were deleted (Figure 4B); the construct Sticky^774-1078^–GFP and smaller truncations were no longer recruited to the cortex. Thus, we conclude that residues 974-1128 are required for the cortical localization of Sticky’s CC2a domain. However, we noted that some of the shorter constructs (Sticky^774-1078^–GFP, Sticky^774-1028^–GFP, Sticky^774-978^–GFP) that did not localize to the cortex nonetheless exhibited a later midbody localization, which might indicate additional interactions within the CC2a region (Figure 4B).

**Figure 4.**
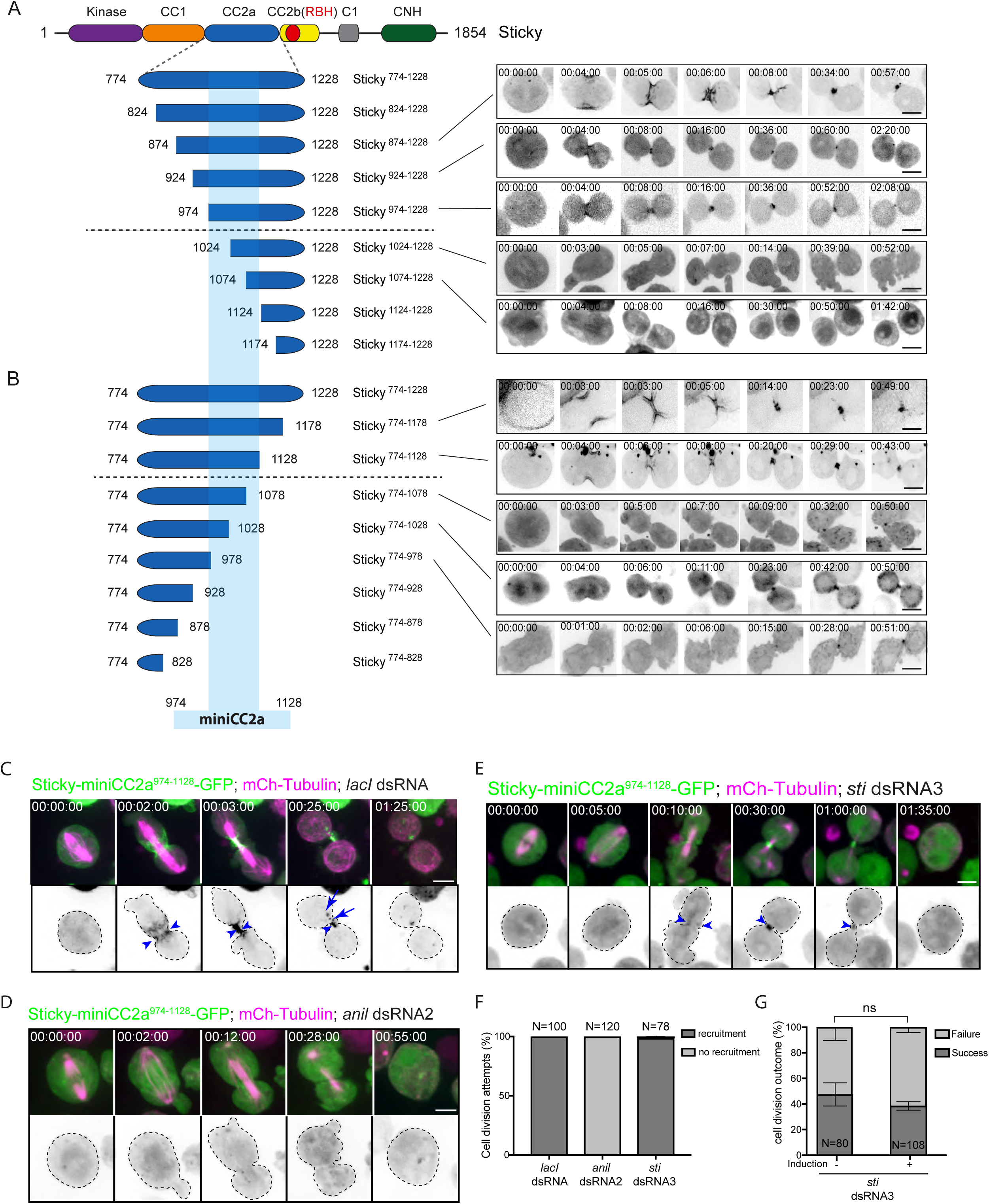
Defining the minimal Anillin-dependent input to Sticky localization. **A-B** Cartoons on the left indicate the series of N-terminal (A) and C-terminal (B) truncations of Sticky-CC2a generated and, on the right, high-resolution time-lapse sequences of the corresponding, representative localization patterns observed for the indicated truncations fused to GFP (inverted grayscale images of the GFP channel are shown). Horizontal black lines separate the cartoon depictions of the constructs that localized to the CR/nascent MR (above the line) from those that did not (below the line), while the light blue shaded region represents the Sticky-miniCC2a^974-1128^ region defined by this truncation analysis. **C-E** Time-lapse sequence of representative cells stably expressing mCherry-Tubulin (magenta) and inducibly Sticky-miniCC2a^974-1128^-GFP (green in upper panel and inverted grayscale in lower panel) incubated for 3 days with control *lacI* dsRNA (C), *anillin* dsRNA2 (D) or sticky dsRNA3 (E). Times are h:min:s; scale bars are 5 µm. **F** Quantification from high-resolution imaging of the percentage of cells that showed cortical recruitment vs. no cortical recruitment of Sticky-miniCC2a^974-1128^-GFP during anaphase/telophase following indicated RNAi treatment. **G** Quantification of the outcome (success or failure) of division attempts scored from low-resolution imaging of cells stably expressing mCherry-Tubulin and induced to express Sticky-miniCC2a^974-1128^-GFP following 3-day incubation with *sti* dsRNA3. Data are from 3 independent experiments. Total number of cells analyzed (N) for each condition is indicated on respective bars. Error bars represent standard deviation between experiments resulting from an unpaired t-test with ns = non-significance.

We next tested whether the residues 974-1128 were sufficient for cortical localization by generating an inducible Sticky^974-1128^-GFP construct and expressing it in S2 cells. This construct, named “miniCC2a”, was robustly recruited to the CR and nascent MR in 100% of control cells observed via live-imaging (Figure 4C, 4F, Supplemental Movie 3, N=100), while 100% of Anillin-depleted cells failed to recruit Sticky-miniCC2a^974-1128^-GFP to the furrow cortex (Figure 4D, 4F, N=120, Supplemental Movie 4). In cells depleted of endogenous Sticky (dsRNA3), Sticky-miniCC2a^974-1128^-GFP was still recruited in 100% of cases (N=78), but failed to rescue cytokinesis since 61% of these division attempts failed (N=108), compared to 56% in the uninduced controls (N=80, Figure 4E-G, *p*=0.467). We noted that in control cells expressing endogenous Sticky, Sticky-miniCC2a^974-1128^-GFP was excluded from mature MRs (e.g. Figure 4C), suggesting that Sticky-miniCC2a^974-1128^-GFP lacks elements required for the formation of the MR and/or its retention at the MR. Given that a Protein A-CC2a^774-1227^ fusion was reported to pull-down actin and myosin from cell extracts (Bassi *et al.*, 2011), we tested for potential contributions of actomyosin in recruiting Sticky-miniCC2a^974-1128^-GFP. Prior studies have shown that, when the actin cytoskeleton is disrupted with Latrunculin A (LatA) which prevents actin polymerization (Coue *et al.*, 1987; Ayscough *et al.*, 1997; Ayscough, 1998), Rho1 drives the assembly of Anillin-and septin-dependent structures at the equatorial membrane during anaphase (Hickson and O’Farrell, 2008b). Persistent cortical localization of Sticky-miniCC2a^974-1128^-GFP in 1 µg/ml LatA, indicates that its recruitment is actin-independent (Supplementary Figure S3). Recruitment of Sticky-miniCC2a^974-1128^-GFP was also unaffected, in the presence or absence of LatA, by depletion of Rho-kinase (Rok; Supplementary Figure S3F), myosin heavy chain (Zipper; Supplementary Figure S3G), or myosin regulatory light chain (Spaghetti squash; data not shown), indicating that recruitment is independent of myosin II. Depletion of kinesin 6/Pavarotti or Kif14/Nebbish, that can also bind to the CC1 region of Sticky during cytokinesis (Bassi *et al.*, 2013), had no effect on Sticky-miniCC2a^974-1128^-GFP recruitment either (Supplemental Figure S3H-I). Indeed, of all candidates tested, only Anillin or RhoGEF/Pebble RNAi diminished cortical recruitment of Sticky-miniCC2a^974-1128^-GFP, which were assayed both in the presence and absence of LatA (Figure 4D, Supplemental Figure S3D-E and data not shown). We therefore conclude that residues 974-1128 of Sticky are sufficient for cortical recruitment to the equatorial cortex in a manner that specifically requires RhoGEF/Pebble and Anillin, but does not require Rho-kinase, actomyosin, kinesin-6/Pav or Kif14/Nebbish.

### Mapping the RBD-dependent contribution to Sticky localization

Our analysis of the L1246N mutation in the context of full-length Sticky (Figures 1D and 2B and Sticky-CC2a-CC2b (Figure 3) indicated the involvement of the RBH domain in Sticky localization. However, Bassi *et al.* (2011) had shown that a construct comprising only residues 1228-1386, which harbors the RBH domain and which they termed CC2b, was not recruited to the cortex. We confirmed this finding using Sticky-CC2b-GFP (residues 1229-1370; Figure 5A), but also generated an additional series of constructs that extended this construct in ∼50-residue increments in the N-terminal direction. These extended constructs now localized to the CR (Figure 5B, D & F), in a manner that was inhibited by inclusion of the L1246N mutation (Figure 5C, E & G), indicating that they depended on a functional RBH. The only constructs that still localized to the CR when harboring the L1246N mutation were the ones that also included the entire Anillin-dependent miniCC2a^974-1128^ residues (Figure 5H-I). Importantly, Sticky residues 1174-1370, which we refer to here as “maxiCC2b”, promoted CR localization without any overlapping amino acid residues with the miniCC2a (residues 974-1128) region, confirming that these two inputs are physically separable, albeit directly adjacent.

**Figure 5.**
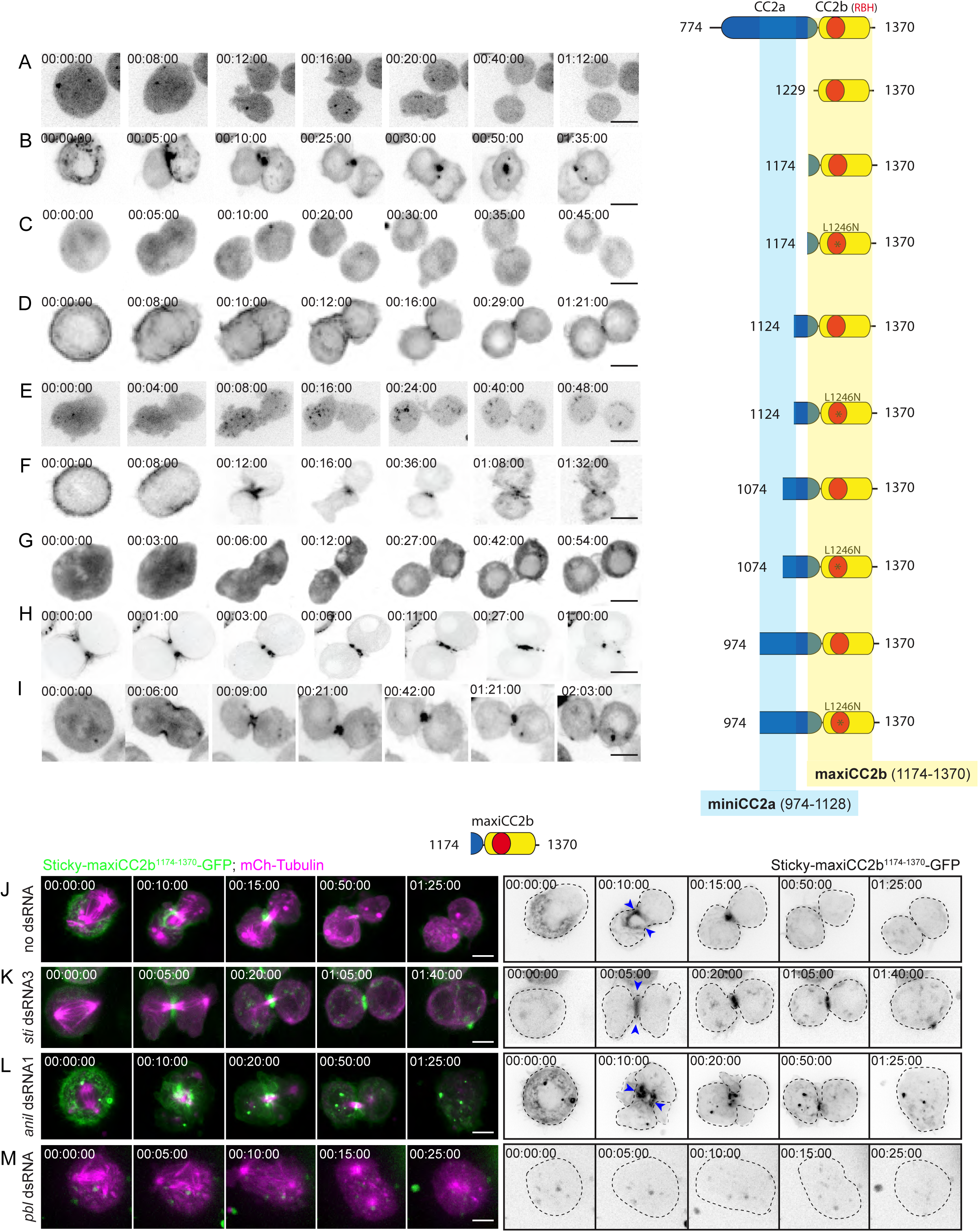
Mapping the RBD-dependent contribution to Sticky localization. **A-I** High-resolution time-lapse sequences of the representative localization patterns observed for the Sticky-CC2a-CC2b-GFP N-terminal truncations and L1246N mutants are shown on the left and depicted in the cartoons on the right. Inverted grayscale confocal images of the GFP channel are shown. Only Sticky-CC2b^1229-1370^GFP failed to localize to the cell cortex (A), whereas the longer GFP constructs were recruited to the cortex (B, D, F, H) in a manner that was abolished by the L1246N mutation (C, E, G), except for when the full miniCC2a^974-1128^ sequence was also present (I). The yellow shaded area of the cartoons represents the minimal RBH-dependent sequence (residues 1174-1370) required for cortical recruitment of maxiCC2b that is distinct from the miniCC2a sequence (residues 974-1128; light blue shading). **J-M** Representative, high-resolution time-lapse sequences of cells transiently expressing mCherry-Tubulin (magenta) and inducibly expressing Sticky-maxiCC2b^1174-1370^-GFP (green in the merged panels on the left; inverted grayscale in the panels on the right) following 3-day incubation with the specified dsRNAs. Blue arrowheads highlight cortical recruitment at the CR and dashed lines indicate the outline of the cell in the grayscale images. Times are h:min:s; scale bars are 5 µm.

We further confirmed that Sticky-maxiCC2b^1174-1370^-GFP (Figure 5J) was recruited independently of endogenous Sticky (Figure 5K) and Anillin (Figure 5L). However, as expected, Pebble RNAi blocked equatorial enrichment of Sticky-maxiCC2b^1174-1370^-GFP (Figure 5M). Collectively, these results support the conclusion that the CC2 region of Sticky contains at least two separable Rho-dependent inputs that contribute to furrow localization: one through residues 974-1128 (miniCC2a; Rho1- and Anillin-dependent), and another one through residues 1174-1370 (maxiCC2b; Rho1-dependent, but Anillin-independent).

### The localization of miniCC2a specifically requires the N-terminal domain of Anillin

We sought to further define how Anillin contributes to Sticky localization. Anillin is a multi-domain, Rho-dependent scaffold protein (Hickson and O’Farrell, 2008b, a; Piekny and Glotzer, 2008). We previously showed that Sticky acts to retain the N-terminal half of Anillin (residues 1-802), which lacks the C-terminal Anillin homology (AH) and Pleckstrin homology (PH) domains, but is sufficient to form a MR-like structure (El Amine *et al.*, 2013). We therefore hypothesized that Anillin would require sequences within its N-terminal half to recruit miniCC2a, even though the Rho-dependent recruitment of Anillin occurs via its C-terminal AH-PH region (Piekny and Glotzer, 2008; Kechad *et al.*, 2012; Sun *et al.*, 2015). To test this hypothesis, we generated stable cell lines co-expressing Sticky-miniCC2a^974-1128^-mCherry and various Anillin truncations fused to GFP (Figure 6A), and examined their localization patterns in live cells depleted of endogenous Anillin. While 100% of cells expressing full-length Anillin-GFP robustly recruited miniCC2a to the contractile ring (Figure 6B and D, N=70), cells expressing Anillin lacking its N-terminal domain (NTD) of 147 amino acids (Anillin-ΔNTD-GFP) failed to recruit miniCC2a in 95% of division attempts (Figure 6C and D, N=49 \*\**p*=0.0032). Upon depletion of endogenous Anillin by RNAi, Anillin-ΔNTD failed to recruit miniCC2a in 100% of division attempts (Figure 6D, N=65 for *anil* dsRNA2 and N=53 for *anil* dsRNA 3’UTR, \*\*\**p*=0.00012). We next wished to test the recruitment of miniCC2a to the Rho1-driven, Anillin- and septin-dependent structures that form at the equatorial membrane during anaphase, upon disruption of the actin cytoskeleton with LatA. (Hickson and O’Farrell, 2008b). Anillin-GFP recruited miniCC2a to these structures (Figure 6E, Supplemental Movie 5) following LatA treatment, while Anillin-ΔNTD-GFP did not (Figure 6F, Supplemental Movie 6), confirming that F-actin is not required for the Anillin-dependent recruitment of miniCC2a. Constructs lacking Anillin’s N-terminal myosin binding (Anillin-ΔMyoBD-GFP) or actin-binding (Anillin-ΔActBD-GFP) domain, were still able to recruit miniCC2a to the cleavage furrow (Figure 6G-H). Thus, the NTD of Anillin is specifically required for the recruitment of miniCC2a to the cleavage furrow, presumably reflecting an interaction between Anillin and Sticky. Consistent with this notion, a fragment of Anillin comprising only the NTD fused to GFP was recruited to the nascent MR in control (*lacI* dsRNA, data not shown) and Anillin-depleted cells (*anil* dsRNA1; Figure 6I), whilst this recruitment was lost following Sticky depletion (*sti* dsRNA2; Figure 6J).

**Figure 6.**
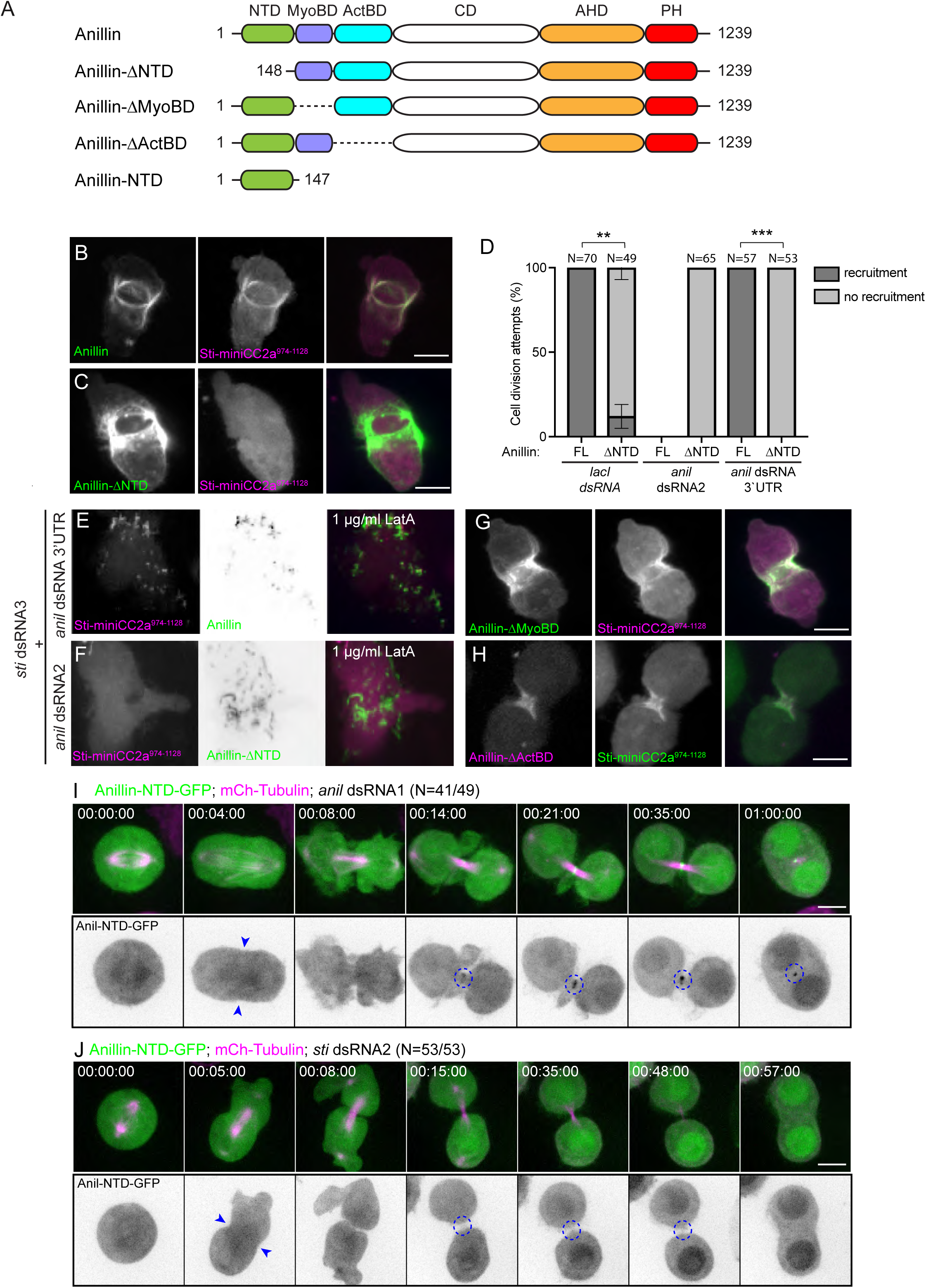
The localization of Sticky-miniCC2a^974-1128^ specifically requires the Anillin N-terminal domain (NTD) **A** A cartoon representation of Anillin showing different domains and truncated constructs. **B-C** Stills from time-lapse sequences of cells mid-anaphase, inducibly expressing Sticky-miniCC2a^974-1128^-mCherry (middle panels, magenta in merged) and (A) Anillin-GFP or (B) Anillin-ΔNTD-GFP (left panels, green in merged). **C** Quantification from time-lapse recordings of the percentage of cells that showed cortical recruitment vs. no cortical recruitment of Sticky-miniCC2a^974-1128^-mCherry during anaphase/telophase in cells co-expressing either Anillin-GFP (FL) or Anillin-ΔNTD (ΔNTD) following 3-day incubation with the specified dsRNAs. N values indicate the total number of cells scored for recruitment per condition and data are from 3 independent experiments. Error bars represent standard deviation between experiments and \*\**p* < 0.01; \*\*\**p* < 0.001 for significance in an unpaired t-test. **D-G** Stills from time-lapse sequences of cells mid-anaphase, inducibly expressing Sticky-miniCC2a^974-1128^mCherry (left panels, magenta in merged) and (D) Anillin-GFP or (E) Anillin-ΔNTD-GFP (grayscale middle panel and green in merged) after a 3-day incubation with *sti* and *anil* dsRNAs as indicated, and following pre-treatment with 1 µg/ml LatA. **F-G** Stills from time-lapse sequences of cells mid-anaphase, inducibly expressing (F) Sticky-miniCC2a^974-1128^-mCherry (middle panel, magenta in merged) and Anillin-ΔMyoBD-GFP (left panel, green in merged) or (G) Sticky-miniCC2a^974-1128^GFP (middle panel, green in merged) and Anillin-ΔActBD-mCherry (left panel, magenta in merged). **H-I** Stills from time-lapse recordings of cells expressing mCherry-Tubulin (magenta in merged upper panels) and Anillin-NTD-GFP (green in upper merged panels; grayscale in lower panels showing inverted images) progressing through cytokinesis following a 3-day incubation with *anil* dsRNA1 (H) or *sti* dsRNA2 (I). Blue arrowheads highlight the ingressing furrow, dashed circles highlight the nascent MR. Times are h:min:s; scale bars are 5 µm.

### A physical interaction between the Anillin NTD and Sticky miniCC2a

Biochemical experiments were employed to test the hypothesis that the Anillin NTD physically interacts with the miniCC2a region of Sticky. GST fusion proteins were expressed and purified from *E. coli* and used in pull-down assays. Unlike GST alone, GST-Anillin-NTD effectively pulled-down endogenous Sticky from untransfected S2 cell lysate (Figure 7A). GST-Anillin-NTD also specifically pulled down miniCC2a-GFP from cell lysate expressing this construct (Figure 7B). Reciprocally, a GST-miniCC2a fusion protein, but not GST alone, effectively pulled down endogenous Anillin from untransfected S2 cell lysate (Figure 7C). Further, GST-miniCC2a pulled down Anillin-NTD-GFP from transfected cell lysate (Figure 7D), but did not pull down Anillin-ΔNTD-GFP (Figure 7E). These data lead to the conclusion that the miniCC2a region (residues 974-1128) of Sticky interacts, directly or indirectly, with the NTD of Anillin (residues 1-147). This supports the observations regarding the recruitment of Sticky-miniCC2a to the CR and nascent MR during live-cell imaging presented in Figures 4 and 6.

**Figure 7.**
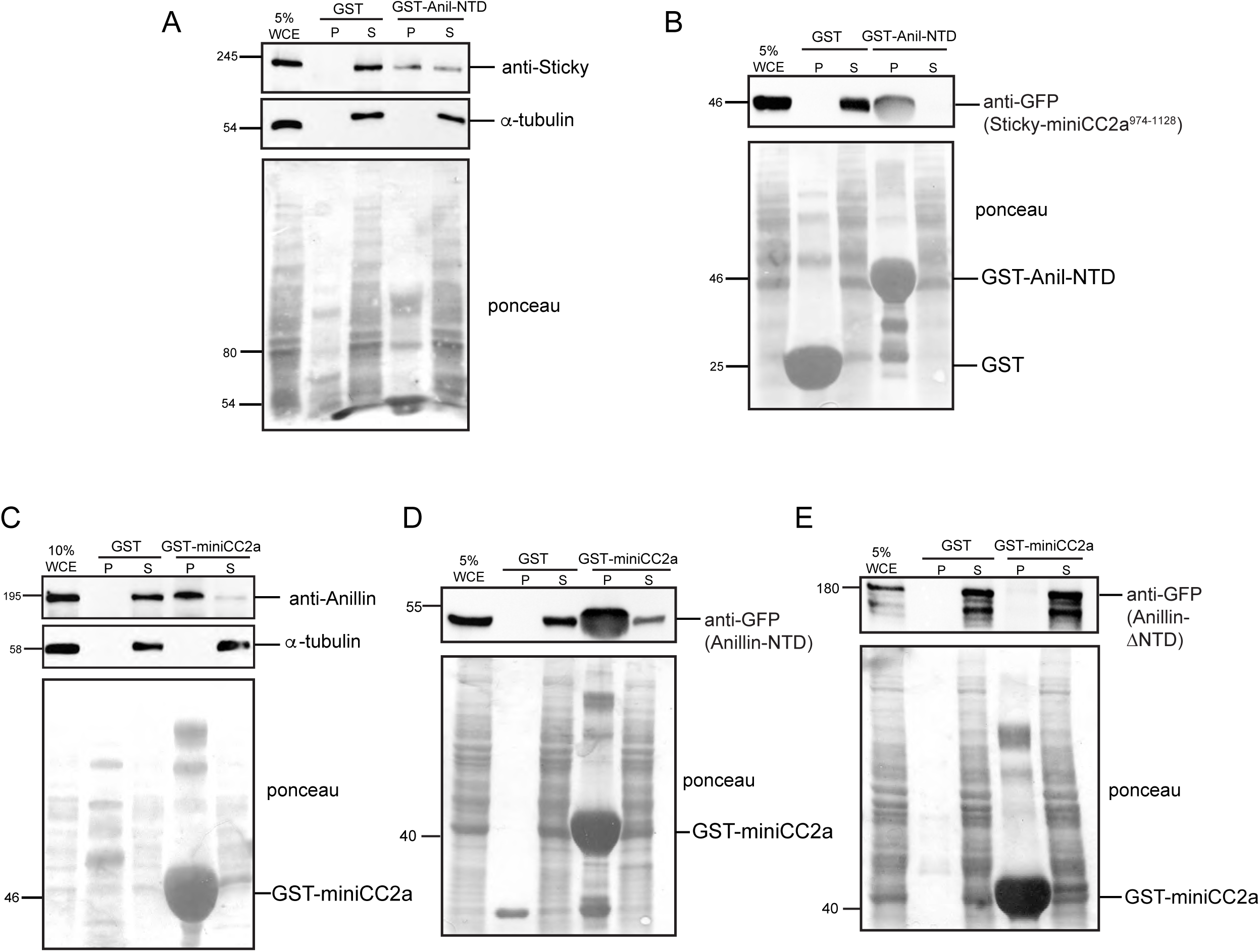
A physical interaction between the Anillin NTD and Sticky-miniCC2a^974-1128^. **A-B** GST-pulldown experiments in which GST-Anillin-NTD was used to pull down endogenous Sticky (A) or Sticky-miniCC2a^974-1128^-GFP (B) from S2 cell whole cell extracts (WCE). Pull-down fractions (P) and supernatants (S) were separated by SDS-PAGE, transferred to membranes and immunoblotted with antibodies against Sticky (A) or GFP (B). **C-E** Analogous blots for which GST-Sticky-miniCC2a^974-1128^ was used to pull down endogenous Anillin (C), Anillin-NTD-GFP (D) or Anillin-ΔNTD-GFP (E) from S2 cell whole cell extracts (WCE). Membranes were immunoblotted with antibodies against Anillin (C) or GFP (D-E). Ponceau S-stained membranes prior to blotting are shown below (A-E) and, in A and C, prior to being cut into two for separate immunoblotting with alpha-Tubulin antibody. The position of GST and GST-fusion proteins are indicated on the Ponceau S-stained membranes.

### The Anillin NTD and Sticky miniCC2a domains are each required for a successful CR-to-MR transition

Given that Anillin-ΔNTD-GFP had failed to recruit (Figure 6C), and interact with (Figure 7E) the miniCC2a fragment of Sticky, we tested the ability of Anillin-ΔNTD-GFP to support cytokinesis in rescue experiments. Stably transfected cells were treated with control dsRNA or one of two different dsRNAs targeting endogenous Anillin, induced to express Anillin-ΔNTD-GFP, and scored for the outcome of division attempt by low-resolution live-imaging 3 days post RNAi. In total, 95% of control depleted cells (*lacI* RNAi) successfully divided (Figure 8A, N=203 for *lacI* dsRNA uninduced cells and N=314 for *lacI* dsRNA induced cells, p=0.714). Conversely, 50% of cells depleted of endogenous Anillin failed cytokinesis (Figure 8A, N=238 for *anil* dsRNA2 induced cells, \*\*\**p*=0.0006 and N=175 for *anil* dsRNA 3′UTR induced cells, \**p*=0.024). This indicates that Anillin requires its 147 amino acid NTD for faithful cytokinesis. High-resolution imaging of these cells also revealed evidence of shedding of Anillin-ΔNTD-GFP (Figure 8B-D, Supplemental Movie 7), although cells often failed cytokinesis with an internal Anillin-ΔNTD-positive structure (Figure 8C-D, Supplemental Movie 7). This suggests that the normal retention of Anillin at the MR does not solely depend on the NTD, and other mechanisms must exist. As an additional test of the role of the Anillin NTD, we examined the behaviors of Anillin-NTD-GFP and Anillin-ΔNTD-mCherry co-expressed in the same cells. This confirmed specific retention of Anillin-NTD at the MR with concomitant shedding of Anillin-ΔNTD (Figure 8E, Supplemental Movie 8).

**Figure 8.**
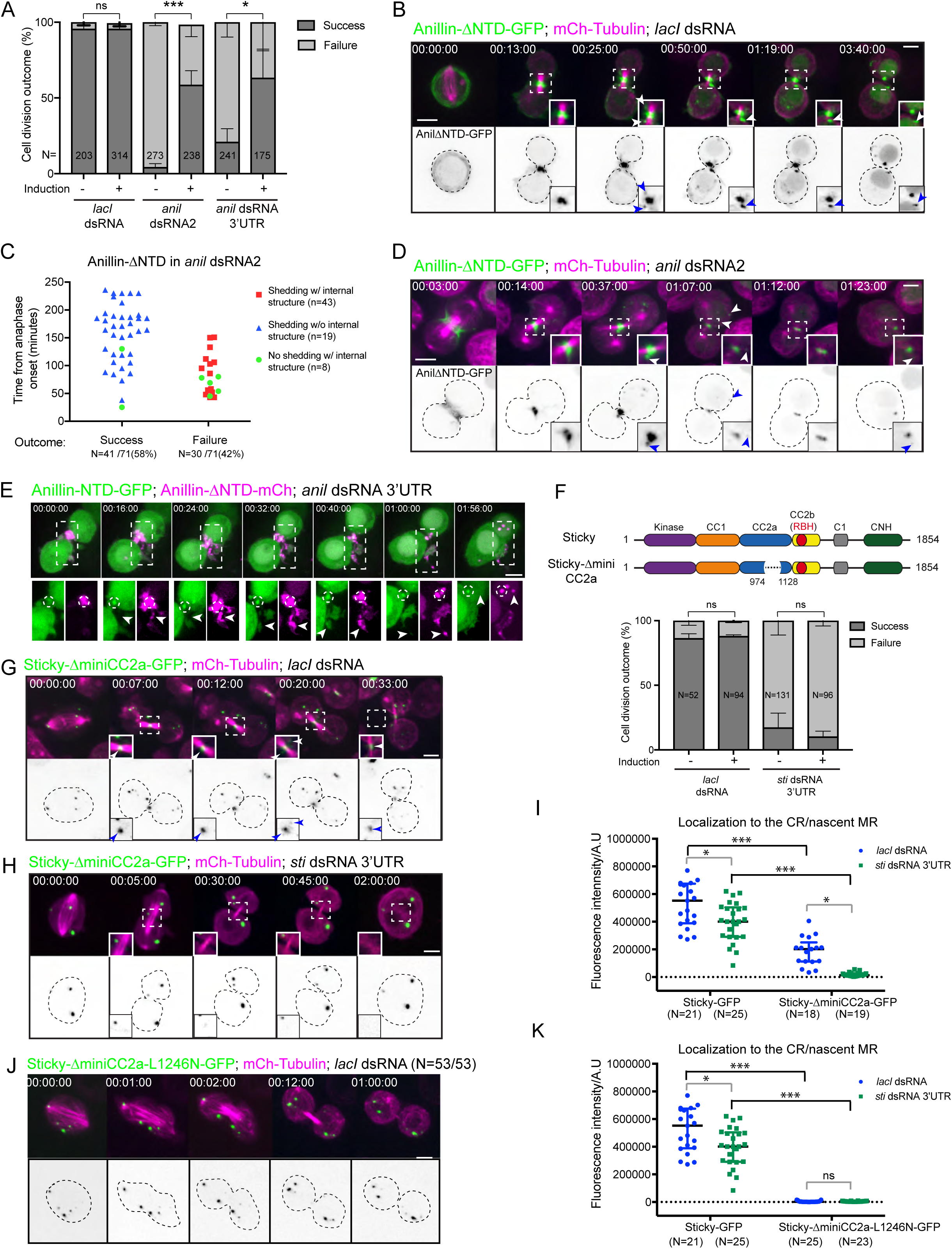
The Anillin NTD and Sticky miniCC2a^974-1128^domains are each required for a successful CR-to-MR transition. **A** Quantification of the outcomes (success or failure) of division attempts in percentage scored from low-resolution time-lapse imaging of cells stably expressing mCherry-Tubulin and inducibly expressing Anillin-ΔNTD-GFP, following 3-day incubation with the specified dsRNAs. N values indicate the total number of cells scored per condition and data are from 3 independent experiments. Error bars represent standard deviation between experiments and * *p*<0.05, \*\*\**p*<0.001 for significance and ns = non-significance in an unpaired t-test. **B**, **D** High-resolution time-lapse sequences of representative examples of the most prevalent phenotypes observed for cells expressing mCherry-Tubulin (magenta) and Anillin-ΔNTD-GFP (green in upper panels, inverted grayscale in lower panels) progressing through cytokinesis, following 3-day incubation with *lacI* (B) or *anil* (D) dsRNAs. Insets are magnifications of the boxed regions and dashed white boxes indicate the region shown in the insets. Blue arrowheads highlight shed particles of Anillin-ΔNTD-GFP and dashed lines indicate the outline of the cell in the grayscale images. **C** Scatter plot of the phenotypes observed by high-resolution imaging of the Anillin RNAi condition. The relative times of success (abscission) or failure (binucleation) of individual cells are plotted and color-coding reflects whether or not shedding of Anillin-ΔNTD-GFP was observed, put in relation to whether or not an internalized Anillin-ΔNTD-GFP-positive structure was observed. **E** High-resolution time-lapse sequence of a cell co-expressing Anillin-ΔNTD-mCherry (magenta) and Anillin-NTD-GFP (green) following a 3-day Anillin RNAi treatment. Lower panels represent separated channels of the dashed boxed region in upper panels. Dashed circles highlight the nascent MR enriched in Anillin-NTD-GFP, arrowheads highlight shed particles enriched in Anillin-ΔNTD-mCherry. **F** Top, a cartoon representation of Sticky and Sticky-ΔminiCC2a. Bottom, quantification of the outcomes (success or failure) of division attempts in percentage scored from low-resolution time-lapse imaging of cells stably expressing mCherry-Tubulin and inducibly expressing Sticky-ΔminiCC2a-GFP, following 3-day incubation with the specified dsRNAs. N values indicate the total number of cells scored per condition and data are from 3 independent experiments. Error bars represent standard deviation between experiments and \*\*\**p*<0.0001 in an unpaired t-test. **G**, **H**, **J** High-resolution time-lapse sequences of the most prevalent phenotypes observed for cells transiently expressing mCherry-Tubulin (magenta) and inducibly expressing Sticky-ΔminiCC2a-GFP (G, H) or Sticky-ΔminiCC2a-L1246N-GFP (J, green in upper panels, inverted grayscale in lower panels) progressing through cytokinesis, following 3-day incubation with *lacI* (G, J) or *sti* (H) dsRNAs. Insets are magnifications of the boxed regions. Arrowheads in G highlight the nascent midbody and shed particles of Sticky-ΔminiCC2a-GFP. Times are h:min:s; scale bars are 5 µm. **I**, **K** Scatter plots of the quantification of localization of Sticky-GFP and Sticky-ΔminiCC2a-GFP (I) or StickyΔminiCC2a-L1246N-GFP (K) to the nascent MR by measuring fluorescence intensity at 20 minutes after anaphase onset for all cells analyzed from high-resolution time-lapse sequences acquired after a 3-day incubation with the specified dsRNAs. N values indicate the total number of cells scored per condition and data are from 3 independent experiments. Data is displayed as median with the interquartile range and \**p≤*0.05, \*\*\**p*<0.0001 for significance and ns=non-significance in a Mann-Whitney U test.

Similar deplete-and-rescue experiments were performed to test the requirement for the miniCC2a domain for the localization and function of Sticky. Cells were induced to express Sticky-ΔminiCC2a-GFP and the outcome of division attempts was scored through low-resolution live imaging. In control *lacI* dsRNA-treated cells, 90% of divisions were successful (Figure 8F, N=52 for *lacI* dsRNA uninduced cells, N=94 for *lacI* dsRNA induced cells, *p*=0.473), while 95% of cells treated for 3 days with Sticky 3’UTR dsRNAs failed cytokinesis (Figure 8F, N=131 for *Sti* dsRNA 3’UTR uninduced cells and N=96 for *Sti* dsRNA 3’UTR induced cells *p*=0.366). Thus, amino acids 974-1128 of Sticky, comprising the miniCC2a sequence, are required for cytokinesis.

High-resolution imaging revealed that in the control *lacI* dsRNA-treated cells, Sticky-ΔminiCC2a-GFP localized to the midbody region after CR closure, although this localization was significantly weaker compared to Sticky-GFP (Figure 8G, I, N=21 for Sticky-GFP and N=18 for Sticky-ΔminiCC2a-GFP, \*\*\**p*≤0.0001). However, in Sticky-depleted cells, Sticky-ΔminiCC2a-GFP localization was still detectable albeit barely. (Figure 8H-I, N=19, *p*≤ 0.0001, Supplemental Movie 9). This suggests that the presence of endogenous, full-length Sticky can contribute to Sticky-ΔminiCC2a-GFP localization. We observed that Sticky-GFP intensity at the MR was also slightly decreased in Sticky-depleted cells (Figure 8I, N=25, *p≤ 0.019). We also introduced the L1246N mutation into Sticky-ΔminiCC2a-GFP, which abolished any localization of this construct at the division site (Figure 8J, N=53), regardless of whether or not endogenous Sticky was depleted (Figure 8K, N=25 for *lacI*-depleted cells and N=23 for *sti*-depleted cells, ***p≤0.0001), This further supports the conclusion that both miniCC2a and the RBH-containing maxiCC2b contribute to Sticky localization, and in a partially redundant fashion although, the dramatic decrease observed in the localization of Sticky-ΔminiCC2a suggests that miniCC2a constitutes the major recruitment mechanism. That Sticky-ΔminiCC2a-GFP was still able to localize to the division site in an L1246N-sensitive manner also indicates that the deletion of miniCC2a did not grossly interfere with the adjacent RBH-containing maxiCC2b region, thereby providing some validation for the use of Sticky-ΔminiCC2a as a separation-of-function allele.

### The Sticky RBD is required for MR maturation

Interfering with the interaction between the Sticky miniCC2a region and the Anillin NTD led to a failure of the CR-to-MR transition. We further explored the consequences of perturbing the RBH-dependent input in the context of full-length Sticky. Despite localizing to the nascent MR, neither Sticky-ΔRBH-GFP nor Sticky-L1246N-GFP was able to rescue for loss of endogenous Sticky (Figure 1G). Upon close inspection of time-lapse sequences, we noted that Sticky-L1246N-GFP was significantly less well recruited to the nascent MR than Sticky-GFP as determined by the lower fluorescence intensity values at the nascent MR exhibited by Sticky-L1246N-GFP compared to Sticky-GFP, irrespective of the presence or not of endogenous Sticky)(Figure 9A, N=26 for *lacI*-depleted cells and N=28 for *sti*-depleted cells, \*\*\**p*≤0.0001 in both cases). Nonetheless, regardless of whether the initial recruitment to the nascent MR was weak or strong, the retention of Sticky-L1246N-GFP at the maturing MR appeared markedly impaired compared to that of Sticky-GFP. The persistence of Sticky-GFP versus Sticky-L1246N-GFP signals at the mature MR was therefore measured over time in high-resolution imaging experiments that were performed in parallel, blind as to the presence or absence of the mutation, and without RNAi (so as not to induce cytokinesis failure). This revealed that while Sticky-GFP signal remained detectable at the mature MR for 249±50 min (mean ± SD., Figure 9B, N=31), Sticky-L1246N-GFP was prematurely lost at 132±56 min (mean ± SD, Figure 9B, N=29, ***p=0.0002). Remarkably, under these same conditions, Sticky-L1246N-GFP also exhibited an increased propensity to be shed than did wild-type Sticky-GFP (Figure 9C-D). Sticky-GFP was only rarely observed to shed from the nascent MR, and in a limited manner (Figure 9C), while Sticky-L1246N-GFP exhibited frequent and extensive shedding (Figure 9D, arrowheads, Supplemental Movie 10). In line with this observation, we also found that the intensities of GFP signal (gray values, Figure 9E) at the maturing midbody (∼ 1 h post anaphase onset) were decreased in cells expressing Sticky-L1246N-GFP (65 AU at peak; corresponding to Figure 9D, t=01:02:00) when compared to those expressing Sticky-GFP (131 AU at peak; corresponding to Figure 9C, t=01:00:00). Enhanced shedding and poor retention at the maturing midbody therefore indicate that the L1246N mutant is compromised in its ability to be incorporated into a stable MR structure that is resistant to shedding. We therefore conclude that the RBH domain of Sticky, which makes only a partially redundant contribution to Sticky recruitment, is required for successful cytokinesis through a role in Sticky retention and stable, mature MR formation.

**Figure 9.**
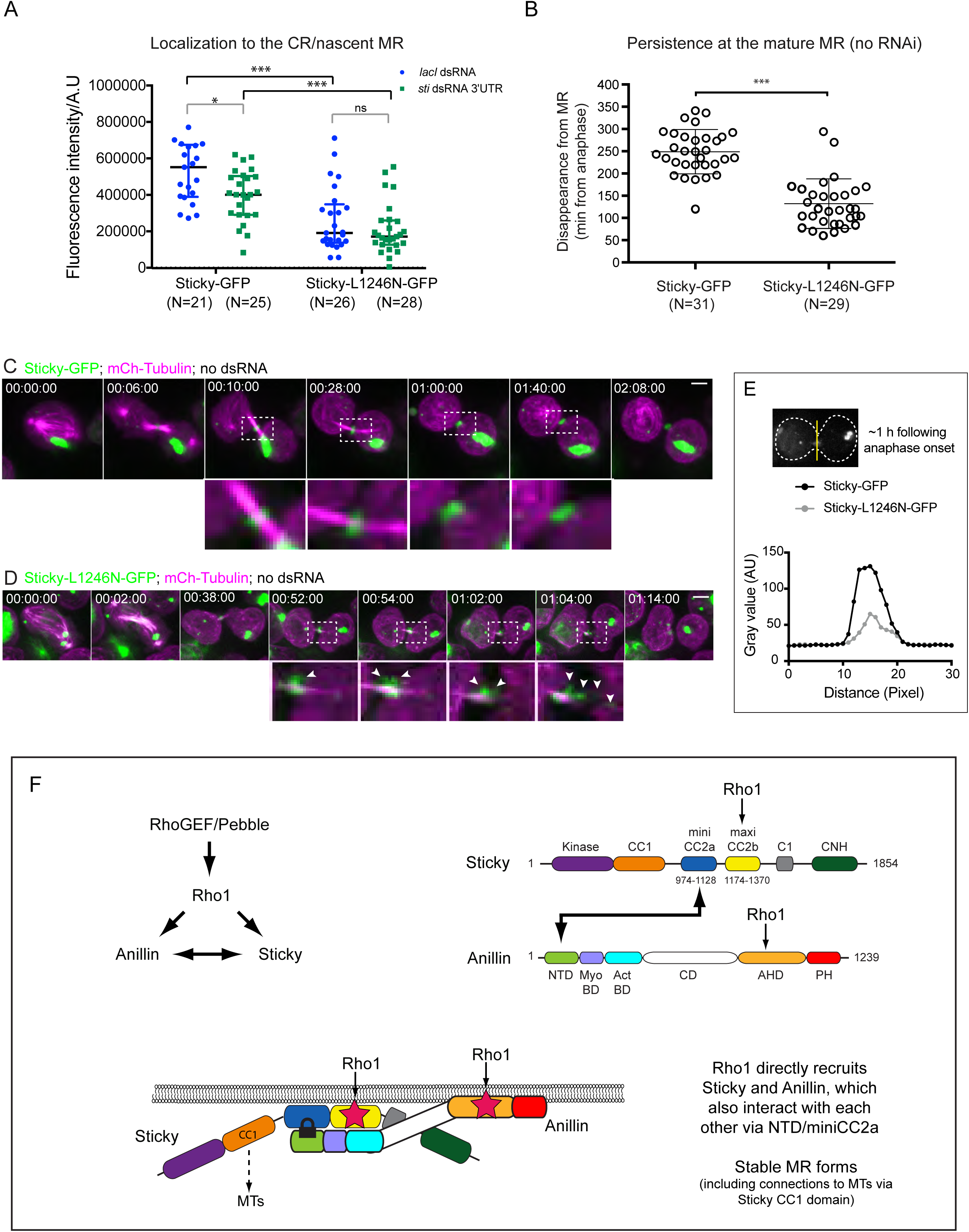
The Sticky RBD promotes its retention at the stable MR. **A** Scatter plot of the quantification of localization of Sticky-GFP and Sticky-L1246N -GFP to the nascent MR by measuring fluorescence intensity at 20 minutes after anaphase onset for all cells analyzed from high-resolution time-lapse sequences acquired after a 3-day incubation with the specified dsRNAs. N values indicate the total number of cells scored per condition and data are from 3 independent experiments. Data is displayed as median with the interquartile range and ***p<0.0001 for significance in a Mann-Whitney U test. **B** Scatter plot of the relative times (in minutes) of disappearance of detectable GFP signal from the mature MR of individual cells expressing Sticky-GFP or Sticky-L1246N-GFP, scored in a blind fashion using high-resolution time-lapse sequences acquired in parallel and without RNAi. N values indicate the total number of cells scored per condition and data are from 2 independent experiments. Error bars represent standard deviation between cells and ***p<0.001 for significance obtained in an unpaired t-test. **C-D** High-resolution time-lapse sequences of cells transiently expressing mCherry-Tubulin (magenta) and inducibly expressing Sticky-GFP (green in C) or Sticky-L1246N-GFP (green in D), progressing through cytokinesis. Lower panels represent magnifications of the dashed boxed regions. Arrowheads highlight shed particles. Times are h:min:s; scale bars are 5 µm. **E** Average line intensity profiles of Sticky-GFP and Sticky-L1246N-GFP across the MR region at ∼1 h post anaphase. Measurements taken for Sticky-GFP correspond to time point 01:00:00 indicated in Fig. 9C, while measurements taken for Sticky-L1246N-GFP correspond to time point 01:02:00 indicated in Fig. 9D. Measurements were taken for one cell per construct. **F** Model showing proposed, cooperative interactions between Rho1, Sticky and Anillin at play during recruitment to the CR/nascent MR and retention at the mature MR.

## Discussion

We report a detailed structure-function analysis that helps clarify the controversial relationship between the Citron kinase, Sticky, and the small GTPase Rho. The work defines two adjacent, Rho-dependent inputs that both contribute to the cortical recruitment and retention of Sticky during MR formation. One input requires interaction with Anillin, itself a Rho-dependent protein (Hickson and O’Farrell, 2008b; Piekny and Glotzer, 2008; Sun *et al.*, 2015), while the other requires the RBH domain of Sticky, presumably via a direct interaction with Rho1-GTP. Although these two inputs are experimentally separable, they function together as a unit with each playing essential roles in ensuring the error-free transition to a stable, mature MR. Overall, the results provide strong indications that Rho, the master regulator of the CR, also controls biogenesis of the subsequent MR through its coordinated actions on Citron kinase and Anillin.

### Interactions of Sticky with Rho

Mammalian Citron kinase was first identified through a yeast two-hybrid screen for RhoA-GTP binding proteins (Madaule *et al.*, 1995; Madaule *et al.*, 1998). It was thus assumed that Citron kinase, and its *Drosophila* ortholog Sticky, would act as canonical Rho effectors: i.e. that direct Rho-GTP binding would drive its recruitment to the equatorial cortex and activate its kinase domain, in much the same way as the related Rho-dependent kinase Rok is thought to be regulated (Di Cunto *et al.*, 1998; Madaule *et al.*, 1998; Madaule *et al.*, 2000; Eda*et al.*, 2001; Shandala *et al.*, 2004; Thumkeo *et al.*, 2013). However, the assumption that Citron kinase/Sticky acts as a canonical Rho-effector at the MR has been called into question on the basis of several observations, as discussed in (D’Avino, 2017). In particular, Gai *et al.* (2011) showed that inhibiting RhoA with cell-permeant C3 exoenzyme during late cytokinesis of HeLa cells led to a loss of Anillin from the intercellular bridge, but not a loss of Citron kinase; whereas depletion of Citron kinase led to a loss of both RhoA and Anillin. This was interpreted to signify that Citron kinase is upstream of RhoA, unlike Anillin, which is downstream of RhoA. Sticky was shown only to bind to Rho1 via its C-terminal CNH domain, and independently of Rho1 nucleotide status (Bassi *et al.*, 2011), while Sticky localization was shown to not require its putative Rho-binding domain (Bassi *et al.*, 2011). Indeed, the structure-function analysis of Bassi *et al.* (2011) failed to identify a role for the RBH domain of Sticky, but the only fragment examined (“CC2b”, residues 1228-1386) lacked essential residues for it to act as a functional Rho-binding domain (RBD). Indeed, the mapping analysis presented here demonstrates that the minimal RBD resides within residues 1174-1370, which we termed maxiCC2b. The present study also clearly shows that the RBD of Sticky is not only functional, in that it contributes to Sticky recruitment to the CR, but also that it is essential, since the Sticky-L1246N mutant predicted to no longer bind Rho-GTP, is unable to form a stable MR that can resist the shedding and disintegration that otherwise occurs. A similar conclusion was reached by Watanabe *et al.* 2013 who found that the coiled-coil domain of mammalian Citron kinase was largely able to rescue the loss of Citron kinase, in a manner that was sensitive to the analogous mutation.

As for the reports of Rho1 interacting with the C-terminal CNH domain of Sticky (Shandala *et al.*, 2004; Bassi *et al.*, 2011), we failed to find any clear functional consequences during cytokinesis for such an interaction, based on deletion of the CNH domain in depletion-and-rescue experiments. This argues that the CNH domain is dispensable for cytokinesis of S2 cells, although it may be important in other cell types.

### A conserved Rho1/Anillin/Sticky complex is required for MR formation

Bassi et al., 2011 identified residues 774-1227 (CC2a) as being sufficient for cortical recruitment of Sticky, and showed that GST-CC2a pulled down actin and myosin from cell lysate. This led to the proposal that the CC2a region localized to the cortex through interactions with actin and myosin filaments. We have further mapped the minimal region required for cortical localization to residues 974-1128 (miniCC2a) and found that its recruitment was uniquely sensitive to Anillin depletion, with persistent recruitment observed upon loss of F-actin (LatA treatment) or myosin II (Rok, MRLC/Sqh or MHC/Zipper depletion). We show that the NTD of Anillin is necessary for the cortical recruitment of Sticky-miniCC2a, in the presence or absence of F-actin (Figure 6), and both necessary and sufficient for the interaction between Anillin and Sticky-miniCC2a (Figure 7). We therefore conclude that the recruitment of Sticky via miniCC2a occurs through interaction with the Anillin NTD, independently of actomyosin. Co-immunoprecipitation of Anillin and Citron kinase from HeLa cells has also been reported (Gai *et al.*, 2011). Despite poor sequence conservation from flies to mammals within the Anillin NTD, truncation analysis also strongly implicated the mammalian Anillin NTD in the interaction. While the interacting region of Citron kinase mapped to more than one region within the C-terminus (Gai *et al.*, 2011), one of these regions (comprising residues 955-1388 of murine Citron kinase) contains the sequence most homologous to Sticky-miniCC2a. Thus, an Anillin-Citron kinase interaction appears to be evolutionarily conserved from flies to mammals. The Anillin-Sticky interaction described here may be direct or indirect, but regardless of the nature of this interaction essential contributions from Rok, myosin II, F-actin, kinesin-6/Pavarotti, KIF14/Nebbish can be excluded on the basis of persistent co-localization of Anillin and miniCC2a following specific depletions and LatA treatment.

Regarding the importance of the Anillin-Sticky interaction to MR formation, we previously showed that depletion of Sticky leads to a complete loss of Anillin at the normal time of MR formation, through mechanisms that include septin-dependent membrane shedding (El Amine *et al.*, 2013). This indicated that Sticky is required to retain Anillin at the mature midbody ring and counteract septin-dependent removal mechanisms. We therefore hypothesized that disrupting the interaction between the Anillin NTD and the Sticky miniCC2a region might lead to a similar phenotype to that observed following Sticky depletion (El Amine *et al.*, 2013). However, in cells expressing endogenous Sticky, deletion of the Anillin NTD, while blocking cytokinesis, often did not result in complete loss of Anillin via shedding (Figure 8C-D). Shedding of Anillin-ΔNTD was frequently observed, but some Anillin-ΔNTD was nonetheless still retained in internal structures after furrow regression (Figure 8D). These were not persistent MR-like structures like those observed upon Anillin-ΔC expression (Kechad *et al.*, 2012; El Amine *et al.*, 2013), but they appeared rather transient and disintegrated over time. However, their existence nevertheless suggests that the interaction between Sticky and Anillin NTD is not the only mechanism through which Anillin is retained during MR formation. Further studies will be required to define these additional mechanisms.

### Rho-dependent control of MR formation

To summarize our findings, we propose a model (Figure 9F and Supplemental Figure S5) in which Pebble-activated Rho1 directly recruits Anillin via its C-terminus, as shown previously (Hickson and O’Farrell, 2008b; Piekny and Glotzer, 2008; Sun *et al.*, 2015), and Sticky via its RBD. At the same time, Anillin and Sticky also facilitate each other’s recruitment via an interaction between their NTD and miniCC2a regions, respectively. That this mutual support works both ways is evident from the observations that Anillin can recruit Sticky-miniCC2a and Sticky-L1246N (this study), while Sticky can recruit Anillin-ΔC (El Amine *et al.*, 2013) and Anillin-NTD (this study). In addition to these two Rho-dependent inputs providing partially redundant contributions to the recruitment of Sticky to the nascent MR, they also appear to be required for Sticky (and therefore Anillin) retention at the mature MR. This is particularly clear for the RBD input, given the observed premature loss of Sticky-L1246N. The two Rho-dependent inputs that we have described, and their shifting importance from recruitment to retention during the transition from CR to MR, can help reconcile the confusion in the literature regarding the relationship between Sticky and Rho. According to the proposed model, Sticky can be considered as both “downstream” of Rho during its recruitment to the CR/ nascent MR, and “upstream” of Rho during its retention at the mature MR. However, in both cases interaction with Rho-GTP (either directly or indirectly via Anillin) is presumably required, so it does not seem unreasonable to consider Sticky as a conventional Rho effector protein.

In considering the overall role of Sticky, it is important to note that the N-terminal portion of the coiled-coil domain (CC1) is also likely essential for MR formation. Constructs expressing the CC1 domain alone localize to the microtubules at the center of the midbody ((Bassi *et al.*, 2011; Bassi *et al.*, 2013) and our unpublished observations). Furthermore, the CC1 domain can bind to microtubule-associated proteins and motors (Bassi *et al.*, 2013; Watanabe *et al.*, 2013), suggesting that it links MR-localized Sticky to the microtubules of the midbody. It will therefore be important in future work to determine how the CC1-dependent activities of Citron kinase are coordinated with the Rho-dependent activities described here to promote formation of a stable MR.

## Materials and Methods

### Molecular biology and cloning

All constructs, except for Sticky-ΔminiCC2a-L1246N were generated by PCR amplification of the ORF, without stop codons, followed by TOPO-cloning into pENTR-D-TOPO (Invitrogen, #K-2400) then by Gateway recombination using LR Clonase into appropriate destination vectors (Drosophila Gateway Vector Collection; T. Murphy, Carnegie Institution for Science, Washington, DC). The *sticky* (CG10522) ORF was PCR amplified from clone RE26327 (*Drosophila* Gene Collection release 2 collection; Drosophila Genomics Resource Center). For all cloning into Gateway destination vectors, initial PCR amplification was carried out with respective forward primers that included the 5’ CACCATG sequence (Appendix 1). The Sticky-L1246N point mutant was generated by site-directed mutagenesis (see Appendix 1 - Primer list). Sticky-ΔRBH lacks the 28 amino acids (residues Δ1235-1263) that form the conserved Rho-binding homology domain, Sticky-ΔCNH lacks the last C-terminal 404 amino acids (residues 1-1450). Sticky-CC2a encodes amino acids 774-1228, Sticky-CC2a-CC2b encodes amino acids 774-1370 and Sticky-miniCC2a encodes amino acids 974-1128. Sticky-ΔminiCC2a encodes amino acids 1-973 and 1129-1854 and therefore lacks amino acids 974-1128 and was generated by PCR by overlap extension followed by cloning into pENTR-D-TOPO and subsequent recombination into pMT-WG. Sticky-ΔminiCC2a-L1246N was generated by first digesting Sticky-L1246N-pENTR and Sticky-ΔminiCC2a-pMT plasmids with MfeI and BsaAI restriction enzymes (New England Biolabs, #R0589 and #R0531) for 2 hours at 37°C followed by treatment of the Sticky-L1246N-pENTR backbone with 1 µl of calf alkaline phosphatase for 30 minutes at 37°C and gel extraction and purification of digested plasmids. The digested plasmid backbone and insert were ligated at 16°C overnight in a ratio of 1:5 followed by transformation into competent *Escherichia coli* cells (DH5α strain) and screening for positive colonies. The remaining Sticky constructs have encoded amino acids indicated in the relevant figure cartoons. All Anillin constructs derive from clone LD23793 (Berkeley Drosophila Genome Project Gold collection; *Drosophila* Genomics Resource Center) that encodes the CG2092-RB polypeptide. Anillin full-length, Anillin-ΔMyoBD, Anillin-ΔActBD have been described previously (El Amine et al., 2013). Anillin-ΔNTD encodes amino acids 148-1239 and lacks the first 147 amino acids whilst Anillin-NTD encodes the first 147 amino acids only. All constructs were verified by sequencing.

### Cell culture and RNAi

Wild-type *Drosophila* S2 cells (UCSF origin) were grown in Schneider’s medium (Invitrogen/Thermo Fisher Scientific, #21720001) supplemented with 10% (v/v) heat-inactivated fetal bovine serum (US origin, Invitrogen/Thermo Fisher Scientific, #16140071), 50 000 units of penicillin and 50 mg/ml Streptomycin at 25°C in ambient CO2. To generate stable cell lines, wild type S2 cells were seeded at a density of 7.5×10^5^ cells/ml and incubated at 25°C overnight before being co-transfected with 1.0 µg of each respective plasmid and 0.25 µg of pCoHygro (Invitrogen) using FuGENE HD (Promega Corporation, #E2311) transfection reagent. Starting from 48 hours after transfection, resistant clones were selected in 0.4 mg/ml Hygromycin B (Invitrogen, #10687010) for 4 weeks. After selection, cells were grown without Hygromycin B in medium supplemented with 10% fetal bovine serum for at least one week prior to use. For transient DNA transfections, wild type S2 cells were plated at a density of 1.0×10^5^ cells/ml a day prior to transfection. 1.0 µg of each relevant plasmid was transfected using FuGENE HD transfection reagent and per reaction, a 3:1 ratio of FuGENE HD:DNA was diluted in serum-free Schneider’s medium in a total volume of 100 μl and incubated at room temperature for 20 minutes, prior to addition to the cells in normal growth medium.

RNA interference in S2 cells was performed using long (250-700 bp) double-stranded RNA (dsRNA). DNA templates were generated in a two-step PCR amplification from cDNA. In the first round of PCR, gene-specific primers (Appendix 1) were used that included a 5’ 8-bp anchor sequence (GGGCGGGT). In a second round PCR, a universal T7 primer containing the anchor (5’-TAATACGACTCACTATAGGGAGACCACGGGCGGGT-3’) was used to generate gene-specific PCR templates that included the T7 promoter sequence at each 5’ end. RNAs were subsequently generated by *in vitro* transcription overnight at 37°C using RiboMAX transcription kits (Promega Corporation). After RQ1 DNAse digestion of the template cDNA, the resulting RNAs were then precipitated in ethanol, resuspended in nuclease-free water, denatured at 95°C and slowly cooled to room temperature to allow the RNA to anneal and form dsRNAs, which were further verified by agarose gel electrophoresis and quantified.

For RNAi experiments, S2 cells were plated in plastic, flat-bottomed 96-well dishes (Falcon) and incubated with 1 µg/ml of relevant dsRNA for 3-6 days (indicated in the figure legends). For analyses after 6-day RNAi, cells were split 1 in 3 into fresh complete medium 3 days after (the first round of dsRNA treatment) followed by addition of 1µg/ml of fresh dsRNA and incubation for a further 3 days. RNAi experiments in stable cell lines were only carried out after the 4-week selection period and in the case of transient transfections, dsRNA incubation was initiated 2 days after transfection. Expression of relevant constructs under control of the metallothionein promoter (pMT plasmids) was induced by the addition of 0.1 mM CuSO4 24 hours prior to imaging, or 20 minutes after the addition of dsRNA, in the case of rescue experiments. One hour prior to live-imaging, cells were transferred into 200 µl of fresh medium in an 8-well chamber slide (Nunc(tm) Lab-Tek(tm) II, Thermo Fisher Scientific). Live-imaging was carried out after 3 days for Sticky, Anillin, Pebble, Pavarotti and Nebbish dsRNAs and after 7 days for Zipper and Rok dsRNA. In the case of co-depletion experiments, these incubation times were maintained, but individual dsRNA incubations in the same experiment were performed in the presence of the LacI negative dsRNA to control for potential effects of combining dsRNAs. For Latrunculin A experiments, Latrunculin A (Calbiochem/EMD Millipore) was gently pipetted to the cells at a final concentration of 1 µg/ml immediately prior to the start of image acquisition.

### Live-cell microscopy

Time-lapse imaging of *Drosophila* S2 cells in Schneider’s medium was performed at room temperature using a spinning-disc confocal system (UltraVIEW Vox; PerkinElmer), comprising a scanning unit (CSU-X1; Yokogawa Corporation of America),a charge-coupled device camera (ORCA-R2; Hamamatsu Photonics), fitted to an inverted microscope (DMI6000 B; Leica Microsystems) and equipped with a motorized piezoelectric stage (Applied Scientific Instrumentation). Image acquisition was controlled using Volocity version 6.3 (PerkinElmer) and was performed in emission discrimination mode. Low-resolution time-lapse imaging was performed using a Plan Apochromat 40x, 0.85 NA air objective with camera binning set to 2 x 2 and 10 µm Z-stacks were acquired with an optical spacing of 1 µm at 4-5 min intervals overnight. High-resolution imaging was performed using a Plan Apochromat 63x oil immersion objectives NA 1.4, with camera binning set to 2 x 2 and 10 µm Z-stacks were acquired with an optical spacing of 0.5 µm at 1-2 minutes interval, unless otherwise stated in the figure legends.

### Immunofluorescence microscopy

Cells expressing Sticky-CC2a-GFP and mCherry-Tubulin were transferred to a 96-well glass-bottomed plate (Whatman) 2 hours before fixation for 5 min in 4% formaldehyde in PBS. After permeabilization and blocking for 1 hour in PBS containing 0.1% Triton X-100 (PTX buffer) and 5% normal goat serum, cells were incubated with primary antibody at 4°C overnight (concentrated mouse anti-Rho1 mAb p1D9 used at 1:1000,), washed with PTX buffer, and incubated for 1 hour with Alexa Fluor 647–conjugated goat anti–mouse antibody (1:500, Molecular Probes) and Hoechst 33258 (1:1000). Cells were washed in PTX and mounted in Fluoromount-G (SouthernBiotech). Images were acquired using the UltraVIEW Vox system described in the previous section using a Plan Apochromat 63x 1.4 NA oil immersion objective without camera binning and in emission discrimination mode.

### Image processing and statistical analysis

At least 3 independent repeats were carried out for each experiment (unless otherwise indicated in the figure legends). Statistical analyses were conducted comparing independent experiments with use of GraphPad Prism 6-7 software (GraphPad Software Inc). Gaussian distribution of data was assessed using the D’Agostino-Pearson test. Means were compared using an unpaired Student’s t-test to analyze data with a normal distribution and medians were compared using the Mann-Whitney U test to analyze data that was not normally distributed. The total number of cells quantified (N value), p values and significance level are indicated in the respective figure legend and in the main text. Fluorescence intensity measurements at the nascent MR were carried out at 20 minutes post-anaphase onset for all cells and all conditions (construct and dsRNA treatment) by drawing a fixed-size ROI around the nascent MR. The average background fluorescence intensity in the green channel at that specific time-point was determined by recording intensity values (A.U) within the fixed-size ROI placed at 3 random positions in the field of view, that excluded the cell being analyzed and also other neighboring cells. The background-subtracted fluorescence intensity value for each cell analyzed per condition is displayed on the scatter plot (Related to Figures 8I, K and Figure 9A). All measurements were made using Volocity Analysis 6.3. Images were processed for publication using Photoshop CS6 (Adobe) or ImageJ (NIH), and assembled as figures using Illustrator CS6 (Adobe). Video files were exported from Volocity as QuickTime videos (Apple).

### Recombinant protein expression

GST-tagged proteins were expressed in BL21 *Escherichia coli* cells. A 5 ml culture of cells was grown overnight in LB with Ampicillin at 37°C with shaking (220 rpm) and used to inoculate 500 ml of fresh LB+Amp and cells were grown at 37°C with shaking until an optical density of 0.6 at A600nm. Recombinant protein expression was then induced by addition of 0.15 mM of isopropyl b-D-1-thiogalactopyranoside (IPTG) and further incubation at 37°C for 4 hours with shaking. Bacterial cells were harvested by centrifugation at 6000g at 4°C followed by resuspension of the pellet in lysis buffer (20 mM HEPES pH 7.4, 0.2 M NaCl, 1 mM EDTA, 5 mM MgCl2, 0.1% Triton X-100, 1mM DTT, 1 mg/ml lysozyme, 0.2 mg/ml Pefabloc), supplemented with complete protease inhibitors (Roche) and lysed at 4°C on ice for 1.5 hours (with intermittent resuspension). Bacterial lysates were then sonicated on ice for 5-10 minutes using a cycle alternating between 30 seconds of pulse and 30 seconds of rest at 50% amplitude. Sonicated lysate was cleared by centrifugation at 12,000g for 20 minutes at 4°C and then applied to pre-washed glutathione sepharose beads (GE healthcare) overnight at 4°C with rotation. GST-tagged proteins bound to beads were separated from the unbound fraction by centrifugation at 4,500 rpm at 4°C for 15 minutes followed by three washes in HMNT lysis buffer (20 mM HEPES, pH 7.4, 10 mM MgCl_2_, 0.1M NaCl, 0.5% Triton X-100).

### Whole cell extract preparation

Wild-type S2 cells or cells expressing the GFP-tagged protein of interest were cultured in T75 culture flasks for 7 days and induced with 0.1 mM of CuSO_4_ 48 hours before harvesting by centrifugation at 1,500 rpm for 5 minutes. Cell pellets were then washed twice in ice-cold PBS prior to lysis in HMNT buffer (500 µl/T75 flask) (20 mM HEPES, pH 7.4, 10 mM MgCl_2_, 0.1M NaCl, 0.5% Triton X-100 and supplemented with a protease inhibitor tablet) by incubating on ice for 45 minutes with vortexing every 5 minutes. Extracts were clarified by centrifugation at 13,200 rpm for 20 minutes at 4°C in a microfuge. For protein knock-down experiments, wild-type S2 cells were grown in 8 wells of a 96-well plate per condition and treated with relevant dsRNA for 3 days before pooling the contents of the 8 wells and harvesting by centrifugation at 1,500 rpm for 5 minutes and extracted in HMNT buffer (100 µl/8 wells) as above. The concentration of protein on beads and whole cell extracts were measured using the BCA assay (Pierce BCA protein assay kit, ThermoFisher Scientific).

### GST pull down assays

S2 whole cell extract (500 µl at 3 mg/ml per pull down) was cleared by two rounds of ultracentrifugation at 55,000 rpm for 15 minutes at 4°C followed by incubation with GST-tagged bait protein, or GST alone, bound to glutathione-Sepharose beads (500 µg/pull down) overnight at 4°C on a rotating platform. After binding, beads were centrifuged at 5,000 rpm for 5 minutes at 4°C to collect the unbound fraction. Beads were then washed 3 times with cold HMNT and the bound fraction was eluted by incubating the beads in 2X SDS sample buffer, boiling at 100°C for 5 minutes and centrifugation at 13,200 rpm for 5 minutes.

### Immunoblotting

Equal amounts of protein from different samples were subjected to SDS-PAGE (6-10% polyacrylamide gels) and transferred to Amersham Protran nitrocellulose membrane (0.45 µm pore size) using constant current (300 mA) for 2-3 hours. Membranes were stained with Ponceau S, digitally scanned, then blocked with 5% milk in PBS containing 0.15% Tween-20 (PBS-T; blocking solution) for 1 hour at room temperature. Where appropriate, membranes were cut at the relevant molecular weight marker and incubated in blocking solution containing primary antibody overnight at 4°C on a rocking platform. Membranes were washed three times for 5 minutes in PBS-T and incubated in horseradish peroxidase-conjugated donkey anti-rabbit, anti-mouse secondary antibody (1:4000, GE Healthcare) or alpaca anti-chicken (1:4000, Immune Biosolutions) in blocking solution for 1 hour at room temperature on a rocking platform. Membranes were then washed 3 times in PBS-T followed by a final wash in PBS before signal development in Clarity Western ECL substrate (Bio-rad laboratories) and visualization on a ChemiDoc MP imager.

### Antibodies

Polyclonal antibodies against *Drosophila melanogaster* Anillin were generated by Immune Biosolutions (Sherbrooke, Quebec, Canada) in chickens against amino acids 299-802 of Anillin fused to GST followed by two rounds of affinity purification of the antibodies, first against the immunogen and then against an N-terminal GST tagged- and C-terminal His tagged-Anillin fusion protein (GST-Anillin^257-298^-His) to remove anti-GST antibodies. The purified antibody was used for immunoblotting at 1:2500 and for validation of the Anillin dsRNAs in Fig. S4. Other polyclonal antibodies used include: rabbit anti-Sticky (used for immunoblotting at 1:3000) raised in rabbits against a fragment of the Sticky protein (amino acids 531–742) fused to a C-terminal 6xHis tag (Paolo D’Avino, University of Cambridge, Cambridge, UK; D’Avino et al., 2004) and rabbit anti-GFP (used for immunoblotting at 1:3000; A-6455, Invitrogen). Monoclonal antibodies used include mouse anti-tubulin antibody (used for immunoblotting at 1:2500; Clone DM1A, Sigma-Aldrich) and concentrated mouse anti-Rho1 antibody (used for immunofluorescence at 1:1000; clone p1D9, Developmental Studies Hybridoma Bank).

## Appendix 1- Primers list

Sticky-455 F: 5’-CACCATGATTTCCGCTACCACCGATGAA-3’

Sticky-774 F: 5’-CACCATGCCTGGATCTTTGACCGAACTG-3’

Sticky-783 R: 5’-AATGGCATTCAGTTCGGTCAA-3’

Sticky-1229 F: 5’-CACCATGTACGTGCAGCGGGACATTAAA-3’

Sticky-1228 R: 5’-GAACTGCTCCTTCTCGGCCAA-3’

Sticky-1370 R: 5’-CGTCGTCTTCAGCTCCACCTG-3’

Sticky-1450 R: 5’-GCTAAGATCATCGGCTGACGG-3’

Sticky-824 F: 5’-CACCATGGAGCAGAGTCTTTCACCCACG-3’

Sticky-874 F: 5’-CACCATGACAGCGAATCTATCGCTCTGG-3’

Sticky-924 F: 5’-CACCATGTCACAGGAGGAAACTCGCCAG-3’

Sticky-974 F: 5’-CACCATGTTGGCCAATGTGCACAGATTA-3’

Sticky-1024 F: 5’-CACCATGGACTCTTGTTTGGTCTTACAG-3’

Sticky-1074 F: 5’-CACCATGCAACTTGATACCCTTCATGAG-3’

Sticky-1124 F: 5’-CACCATGCTCAAGGAGCAGCAGAAGAAG-3’

Sticky-1174 F: 5’-CACCATGGTTAGTTTGAAGGAGGAAAAT-3’

Sticky-828 R: 5’-TGAAAGACTCTGCTCGTTGAA-3’

Sticky-878 R: 5’-CGATAGATTCGCTGTACGCGC-3’

Sticky-928 R: 5’-AGTTTCCTCCTGTGACGTCTT-3’

Sticky-978 R: 5’-GTGCACATTGGCCAAGTGCTC-3’

Sticky-1028 R: 5’-GACCAAACAAGAGTCATTCGC-3’

Sticky-1078 R: 5’-AAATCTCAACTTGATACCCTT-3’

Sticky-1128 R: 5’-CTGCTGCTCCTTGAGATTGAG-3’

Sticky-1178 R: 5’-CTCCTTCAAACTAACCATTTC-3’

Sticky-L1246N F: 5’-GCGCAGCACAAAAAGaacATTGACTACCTTCAG-3’

Sticky-L1246N R: 5’-CTGAAGGTAGTCAATgttCTTTTTGTGCTGCGC-3’

Sticky-ΔminiCC2a F: 5’-ATAATCGAGCACAAGAAGCTGGTGGCGCAGCAG-3’

Sticky-ΔminiCC2a R: 5’-CACCAGCTTCTTGTGCTCGATTATCTCCGACTT-3’

Sticky-ΔRBH F: 5’-CAGCGGGACATTACATTGGCGGACAAACTTTTC-3’

Sticky-ΔRBH R: 5’-GTCCGCCAATGTAATGTCCCGCTGCACGTAGAA-3’

Sticky-ΔCNH F: 5’-CACCATGCCACCCAAGATGGAGCCG-3’

Sticky-ΔCNH R: 5’-GCTAAGATCATCGGCTGACGG-3’

<u>Primers used to generate DNA templates to produce dsRNA.</u> (*Note that the 5*’ *8 bp sequence GGGCGGGT is an anchor sequence allowing for a second PCR amplification using a universal T7 primer, sequence* 5’-TAATACGACTCACTATAGGGAGACCACGGGCGGGT-3’).

*lacI* dsRNA F: 5’-GGGCGGGTTGGTGGTGTCGATGGTAGAA-3’

*lacI* dsRNA R: 5’-GGGCGGGTCGGTATCGTCGTATCCCACT-3’

pbl dsRNA F: 5’-GGGCGGGTATGGAAATGGAGACCATTGAA-3’

pbl dsRNA R: 5’-GGGCGGGTCAGATAGTCCATCTCATTGGC-3’

*sti* dsRNA1 F: 5’-GGGCGGGTGTGAAACCGTTGGTGATATGC-3’

*sti* dsRNA1 R: 5’-GGGCGGGTTTCAACCTCTGGAAGTTATCG-3’

*sti* dsRNA2 F: 5’-GGGCGGGTATGGAGCCGATTAGCGTGCGC-3’

*sti* dsRNA2 R: 5’-GGGCGGGTCTGTGAGCTCGGCCAGATAGA-3’

*sti* dsRNA3 F: 5’-GGGCGGGTGTCACATCCATATTCTTTACG-3’

*sti* dsRNA3 R: 5’-GGGCGGGTCGAGCCCAAAGTCACAGAAGT-3’

*sti* dsRNA 3’UTR F: 5’-GGGCGGGTGCATATAGATGAAGGATATAT-3’

*sti* dsRNA 3’UTR R: 5’-GGGCGGGTTTTAAAATATTATGCGACTTC-3’

*anil* dsRNA1 F: 5’-GGGCGGGTTAGAAATCTATGGCATGTTGGC-3’

*anil* dsRNA1 R: 5’-GGGCGGGTGAGAAAACTGTTAACAACCCGC-3’

*anil* dsRNA2 F: 5’-GGGCGGGTATGGACCCGTTTACTCAGC-3’

*anil* dsRNA2 R: 5’-GGGCGGGTTGGTTCCGCCTCCAGCAGGG-3’

*anil* dsRNA3 F: 5’-GGGCGGGTATGGACCCGTTTACTCAGC-3’

*anil* dsRNA3 R: 5’-GGGCGGGTTCGACTGGACAAATGCGGTTTC-3’

*anil* dsRNA 3’UTR F: 5’-GGGCGGGTGGAACCACCCACTGACCCGCT-3’

*anil* dsRNA 3’UTR R: 5’-GGGCGGGTGCGAGTCATCCTAAATTAAATG-3’

rok dsRNA F: 5’-GGGCGGGTGAACGCCAACCGATCCGATGC-3’

rok dsRNA R: 5’-GGGCGGGTGATACTCGCTTCTGGTACACG-3’

zipper dsRNA F: 5’-GGGCGGGTCCTAAAGCCACTGACAAGACG-3’

zipper dsRNA R: 5’-GGGCGGGTCGGTACAAGTTCGAGTCAAGC-3’

pavarotti dsRNA F: 5’-GGGCGGGTAAATCCGTAACGAAACTAACCG-3’

pavarotti dsRNA R: 5’-GGGCGGGTACAACTGCTCTTGGCAGATACC-3’

nebbish dsRNA F: 5’-GGGCGGGTAGCAACTTCGCTCCGAATTGG-3’

nebbish dsRNA R: 5’-GGGCGGGTCAGGAGACCAAATTCATCTGG-3’

T7 anchor primer: 5’-TAATACGACTCACTATAGGGAGACCACGGGCGGGT-3’

## Acknowledgements

We thank past and present members of the Hickson lab for many useful discussions, and Dr. Jean-Claude Labbé for critical reading of the manuscript. We thank Silvana Jananji for purifying the proteins used to generate the Anillin antibody. We thank Dr. Paolo D’Avino for providing the rabbit anti-Sticky antibody and Dr. Emile Levy for access to a ChemiDoc system. N.E. thanks the *Fondation CHU Sainte-Justine* and the *Dept. Pathologie et Biologie Cellulaire, Université de Montréal* for graduate studentships. GRXH thanks the *Fonds de Recherche du Québec, Santé* (*FRQS*) for research scholarships. This work was supported by grants from Canadian Institutes for Health Research (CIHR) (MOP-97788), the Natural Sciences and Engineering Research Council (NSERC) (RGPIN-2014-05083), Canadian Fund for Innovation (CFI), and the Cole Foundation to GRXH. The authors declare no competing financial interests.

## Author Contributions

G. Hickson and N. El-amine conceived the project. N. El-amine generated initial constructs, performed initial mapping of the CC2a/CC2b region of Sticky and performed preliminary GST-pulldown experiments. S. Carim generated Sticky-miniCC2a and Sticky-ΔminiCC2a, performed and analyzed experiments involving these constructs as well as those of Anillin-ΔNTD, and performed the GST-Anillin-NTD and Sticky-miniCC2a pull-down experiments. D. Wernike confirmed and extended the CC2a/CC2b mapping analysis, generated all L1246N constructs and performed and analyzed experiments involving this mutation, as well as those of Anillin-NTD-GFP. G. Hickson supervised the project. S. Carim and D. Wernike prepared the figures. All authors contributed to writing the manuscript.

