## Supplemental figures for "Rho-dependent control of the Citron kinase, Sticky, drives midbody ring maturation"

### Supplemental Material

Figure S1. **Sticky is efficiently depleted by RNAi**

**A** On top, a cartoon representation of Sticky shows the different domains and underneath, *sticky* mRNA showing the regions targeted by the different dsRNAs used to knock-down Sticky. **B** Western blot showing the efficiency of Sticky depletion after incubation with control *lacI* and *sti* dsRNAs 1-3 and -3'UTR for 3 days, after which whole cell extracts were collected, separated by SDS-PAGE, transferred onto a membrane and immunoblotted with antibodies against Sticky and alpha-Tubulin (loading control). Increasing masses of WCE were loaded for each dsRNA treatment.

Figure S2. **The Sticky CNH domain is not required for cortical localization of Sticky and is not essential for cytokinesis**

**A**, A cartoon representation shows the different domains of Sticky and Sticky-ΔCNH. **B-E** Representative, high-resolution time-lapse sequences of cells stably expressing mCherry-Tubulin (magenta) and inducibly expressing (in green in the merged upper panels, or inverted grayscale in the lower panels) Sticky-ΔCNH-GFP (B-C) or Sticky-L1246N-ΔCNH-GFP (D-E) following 3-day incubation with *sti* dsRNA3 (B, D) or *pbl* dsRNA (C, E). N represents the number of cells displaying similar patterns of GFP localization at the division plane at t=00:08:00-00:10:00/total number of cells scored from 2 independent experiments. Blue arrowheads highlight the ingressing cleavage furrow. Times are in h:min:s and scale bars are 5 µm. **F** Quantification of the outcome (success or failure) of division attempts in percentage scored from low-resolution imaging of cells stably expressing mCherry-Tubulin and induced to express the specified GFP-tagged rescue constructs, following 3-day incubation with *sti* dsRNA3. N represents the number of cells scored for each condition and data are from 2 independent experiments. Error bars represent standard deviation between experiments and ns=non-significance resulting from an unpaired t-test.

Figure S3. **Cortical localization of Sticky-miniCC2a$^{974\text{-}1128}$ is Rho1- and Anillin-dependent but independent of actomyosin and of Pavarotti and Nebbish**

**A** A cartoon representation of Sticky shows the different domains and Sticky-miniCC2a$^{974\text{-}1128}$ construct. **B-I** High-resolution time-lapse sequences of representative S2 cells stably expressing mCh-Tubulin (magenta) and inducibly expressing Sticky-miniCC2a$^{974\text{-}1128}$-GFP (in green in the merged upper panels, or inverted grayscale in the lower panels) following incubation with control *lacI* (B) or *sti* (C) or *anil* (D) or *pbl* (E) or *pavarotti* (H) or *nebbish* (I) dsRNAs for 3 days, or *rok* (F) or *zipper* (G) dsRNAs for 7 days and pre-treatment with 1 µg/ml LatA. N represents the number of cells displaying GFP localization to LatA structures at the division plane/total number of cells scored from 3 independent experiments. Times are shown in h:min:s and scale bars are 5 µm.

Figure S4. **Validation of Anillin antibody and efficiency of Anillin depletion by RNAi**

**A** On top, a cartoon representation of Anillin showing the different domains and underneath, *anillin* mRNA showing the regions targeted by the different dsRNAs used to knock-down Anillin. **B** Western blot showing the efficiency of anillin depletion after incubation with control *lacI* dsRNA and *anillin* dsRNAs 1-3 and 3-3'UTR for 3 days, after which whole cell extracts were collected, separated by SDS-PAGE, transferred onto a membrane and immunoblotted with the generated antibody against Anillin and alpha-Tubulin (loading control). Increasing masses of WCE were loaded for each dsRNA. Ponceau S-stained membranes prior to being cut into two for separate immunoblotting with Anillin and alpha-Tubulin antibodies are shown below. **C** Western blot showing the depletion efficiency of Anillin using control *lacI* dsRNA and *anillin* dsRNAs (dsRNA1 or 3-3'UTR) in cell extracts from wild-type S2 cells (left) and uninduced or uninduced Anillin-GFP expressing cells (right). 20 μg of cell extract was loaded in each condition. Blot also shows recognition of endogenously and exogenously-expressed Anillin by the generated anti-Anillin antibody. Ponceau S-stained membrane prior to blotting is shown below.

Figure S5. **Cartoon representations of some of the experiments performed, depicting proposed interactions between Rho1/Sticky and Anillin and the outcomes at the CR and MR**

**Supplementary movie legends**

Movie 1. **Sticky-GFP is robustly recruited to the CR and MR and exhibits some shedding from the nascent MR.** Time-lapse spinning disc confocal microscopy images of *Drosophila* S2 cell stably expressing Sticky-GFP (green) and mCh-Tubulin (magenta) after treatment with control *lacI* dsRNA for 3 days. Frames were taken every 3 minutes for 3 hours with a 63x, 1.4 NA objective. A maximum intensity z-projection (extended focus) is shown and video is playing at 5 fps. Video corresponds to Figure 1A. Time is shown in h:min:s. Scale bar is 5 μm.

Movie 2. **Depletion of Anillin prevents the cortical recruitment of Sticky-L1246N-GFP** Time-lapse spinning disc confocal microscopy images of *Drosophila* S2 cell stably expressing Sticky-L1246N-GFP (green) and mCh-Tubulin (magenta) after treatment with control *anillin* dsRNA1 for 3 days. Frames were taken every minute 3 for 3 hours with a 63x, 1.4 NA objective. A maximum intensity z-projection (extended focus) is shown and video is playing at 5 fps. Video corresponds to Figure 2B. Time is shown in h:min:s. Scale bar is 5 μm.

Movie 3. **Sticky-miniCC2a$^{974\text{-}1128}$-GFP is robustly recruited to the CR and MR and exhibits shedding from the MR.** Time-lapse spinning disc confocal microscopy images of *Drosophila* S2 cell stably expressing Sticky-miniCC2a$^{974\text{-}1128}$-GFP (green) and mCh-Tubulin (magenta) after treatment with control *lacI* dsRNA for 3 days. Frames were taken every 2.4 minutes for 3 hours and 50 minutes with a 63x, 1.4 NA objective. A maximum intensity z-projection (extended focus) is shown and video is playing at 6 fps. Video corresponds to Figure 4C. Time is shown in h:min:s. Scale bar is 5 μm.

Movie 4. **Sticky-miniCC2a$^{974\text{-}1128}$-GFP fails to be recruited to the CR and MR upon Anillin depletion.** Time-lapse spinning disc confocal microscopy images of *Drosophila* S2 cell stably expressing Sticky-miniCC2a$^{974\text{-}1128}$-GFP (green) and mCh-Tubulin (magenta) after treatment with *anil* dsRNA2 for 3 days. Frames were taken every 1.8 minutes for 3 hours with a 63x, 1.4 NA objective. A maximum intensity z-projection (extended focus) is shown and video is playing at 5 fps. Video corresponds to Figure 4D. Time is shown in h:min:s and scale bar is 5 μm.

Movie 5. **Anillin-GFP recruits Sticky-miniCC2a$^{974-1128}$-mCherry to LatA-induced structures.** Time-lapse spinning disc confocal microscopy images of *Drosophila* S2 cell stably expressing Anillin-GFP (green) and Sticky-miniCC2a$^{974-1128}$-mCherry (magenta) treated with *anil* dsRNA 3'UTR and *sti* dsRNA3 for 3 days, followed by treatment with 1 μg/ml of LatA. Frames were taken every minute for 2 hours and 10 minutes with a 63x, 1.4 NA objective. A maximum intensity z-projection (extended focus) is shown and video is playing at 5 fps. Video corresponds to Figure 6D. Time is shown in h:min:s and scale bar is 5 μm.

Movie 6. **Anillin-ΔNTD-GFP fails to recruit Sticky-miniCC2a$^{974-1128}$-mCherry to LatA-induced structures.** Time-lapse spinning disc confocal microscopy images of *Drosophila* S2 cell stably expressing Anillin-ΔNTD-GFP (green) and Sticky-miniCC2a$^{974-1128}$-mCherry (magenta) treated with *anil* dsRNA 2 and *sti* dsRNA3 for 3 days, followed by treatment with 1 μg/ml of LatA. Frames were taken every minute for 1 hour and 40 minutes with a 63x, 1.4 NA objective. A maximum intensity z-projection (extended focus) is shown and video is playing at 5 fps. Video corresponds to stills shown in Figure 6E. Images were rotated by 90˚ clockwise. Time is shown in h:min:s and scale bar is 5 μm.

Movie 7. **Partial shedding from the MR and subsequent internalization of Anillin-ΔNTD-GFP in a cell that fails cytokinesis after depletion of Anillin.** Time-lapse spinning disc confocal microscopy images of *Drosophila* S2 cell stably expressing Anillin-ΔNTD-GFP (green) and mCh-Tubulin (magenta) treated with *anil* dsRNA2 for 3 days. Frames were taken every 1.6 minutes for 2 hours and 56 minutes with a 63x, 1.4 NA objective. A maximum intensity z- projection (extended focus) is shown and video is playing at 5 fps. Video corresponds to Figure 8D. Time is shown in h:min:s and scale bar is 5 μm.

Movie 8. **Retention of Anillin-NTD-GFP at the MR and shedding of Anillin-ΔNTD-mCherry after depletion of Anillin.** Time-lapse spinning disc confocal microscopy images of *Drosophila* S2 cell stably expressing Anillin-NTD-GFP (green) and Anillin-ΔNTD-mCherry (magenta) treated with *anil* dsRNA 3'UTR for 3 days. Frames were taken every 4 minutes for 3 hours with a 63x, 1.4 NA objective. A maximum intensity z-projection (extended focus) is shown and video is playing at 5 fps. Video corresponds to Figure 8E. Time is shown in h:min:s and scale bar is 5 μm.

Movie 9. **Sticky-ΔminiCC2a-GFP is weakly localized to the late CR/nascent MR after depletion of Sticky.** Time-lapse spinning disc confocal microscopy images of *Drosophila* S2 cell transiently expressing Sticky-ΔminiCC2a-GFP (Δ974-1128; green) and mCh-Tubulin (magenta) treated with *sti* dsRNA 3'UTR for 3 days. Frames were taken every 5 minutes for 3 hours with a 63x, 1.4 NA objective. A maximum intensity z-projection (extended focus) is shown and video is playing at 4 fps. Video corresponds to Figure 8H. Time is shown in h:min:s and scale bar is 5 μm

Movie 10. **Sticky-L1246N-GFP is poorly retained and shed from the nascent MR.** Time-lapse spinning disc confocal microscopy images of *Drosophila* S2 cell transiently expressing Sticky-L1246N-GFP (green) and mCh-Tubulin (magenta). Frames were taken every 4 minutes for 3 hours with a 63x, 1.4 NA objective. A maximum intensity z-projection (extended focus) is shown and video is playing at 5 fps. Video corresponds to Figure 9B. Time is shown in h:min:s and scale bar is 5 µm.

Fig. S1

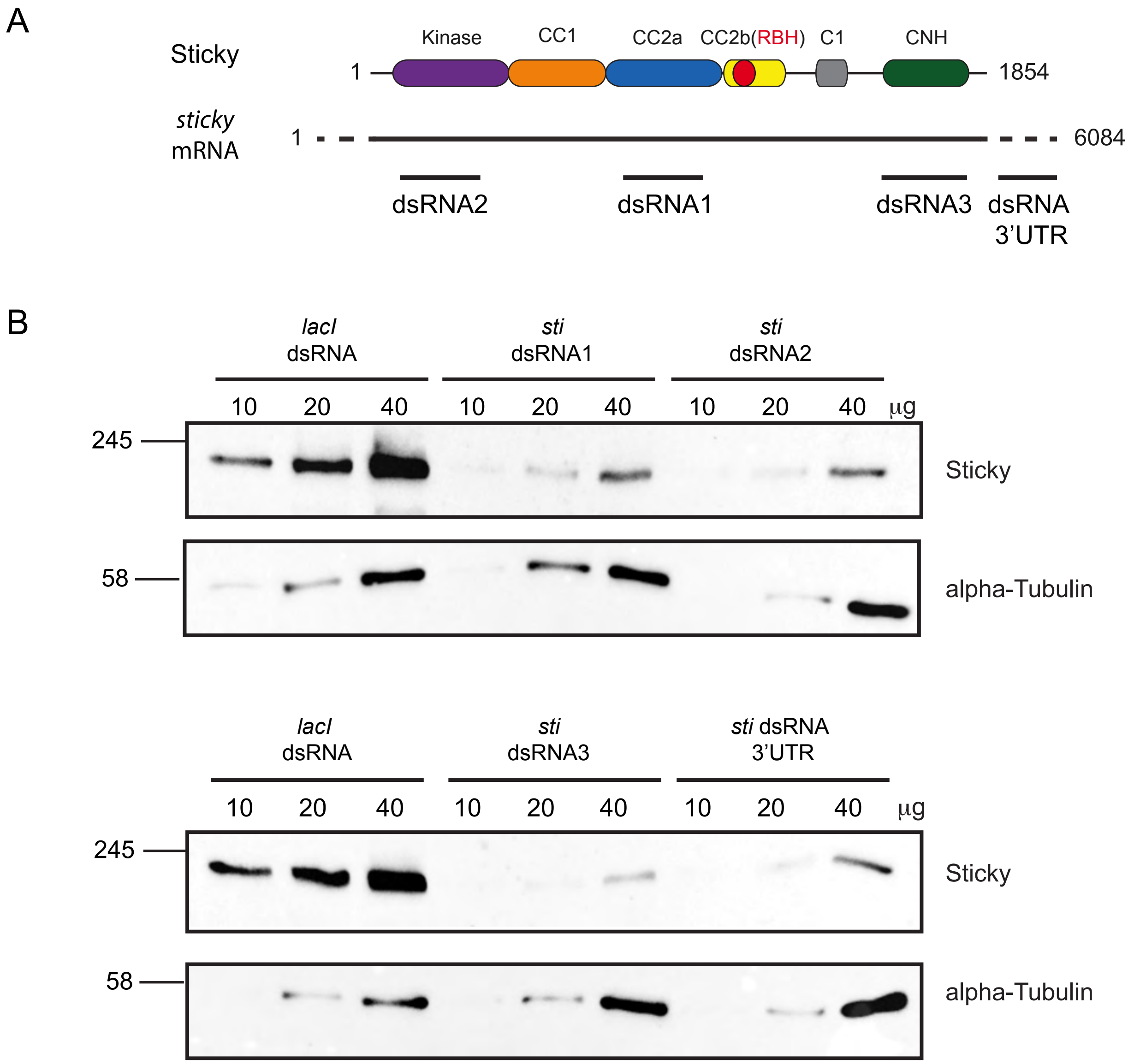

A
Sticky
Kinase
CC1
CC2a
CC2b(RBH)
C1
CNH
1
1854
sticky
mRNA
1
6084
dsRNA2
dsRNA1
dsRNA3
dsRNA
3'UTR
B
lacI
dsRNA
sti
dsRNA1
sti
dsRNA2
10 20 40 10 20 40 10 20 40 μg
245
Sticky
58
alpha-Tubulin
lacI
dsRNA
sti
dsRNA3
sti dsRNA
3'UTR
10 20 40 10 20 40 10 20 40 μg
245
Sticky
58
alpha-Tubulin

Fig. S2

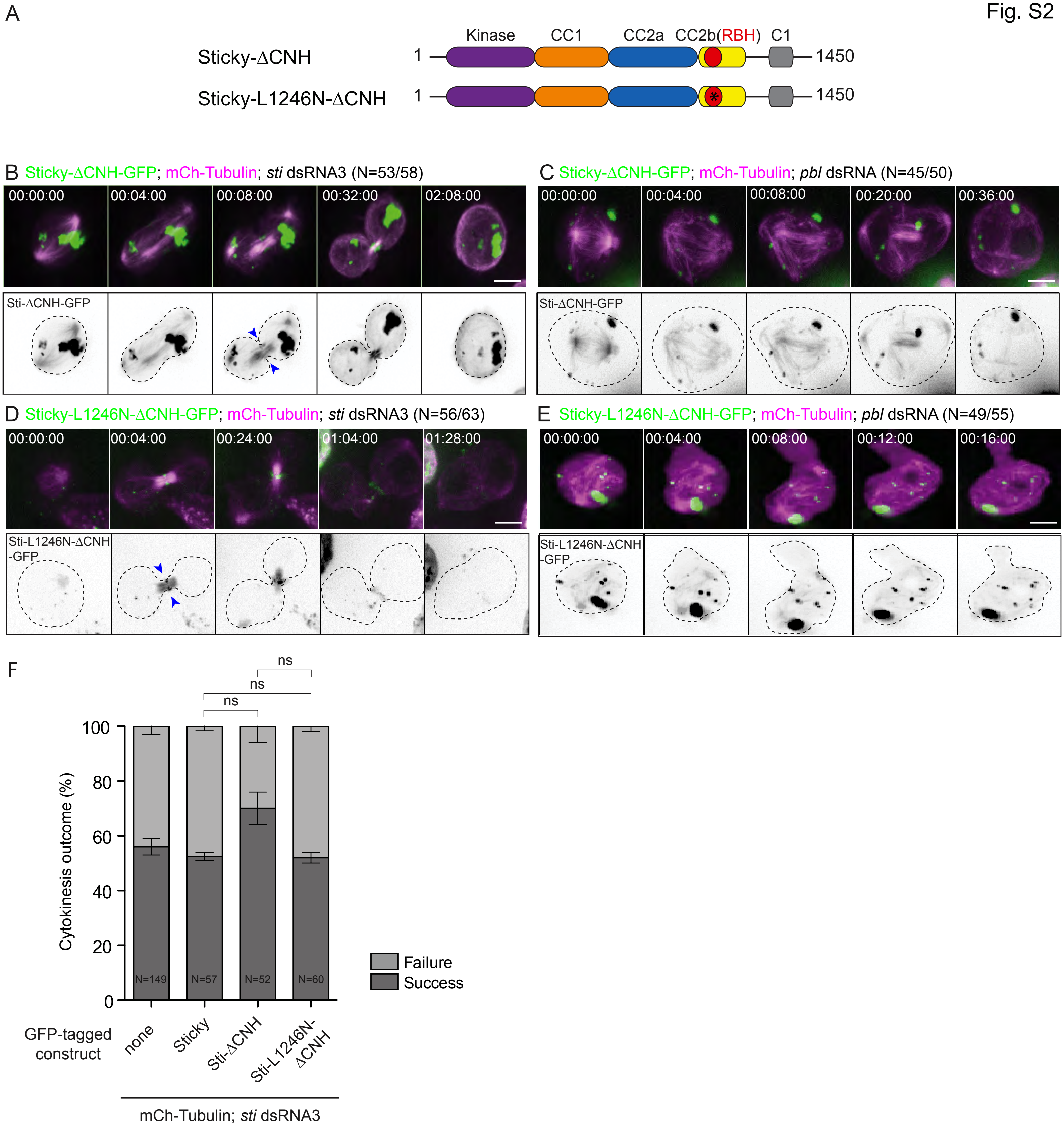

A
Sticky-ΔCNH
Sticky-L1246N-ΔCNH
Kinase
CC1
CC2a
CC2b(RBH)
C1
1
1450
1
1450
B Sticky-ΔCNH-GFP; mCh-Tubulin; sti dsRNA3 (N=53/58)
00:00:00
00:04:00
00:08:00
00:32:00
02:08:00
Sti-ΔCNH-GFP
C Sticky-ΔCNH-GFP; mCh-Tubulin; pbl dsRNA (N=45/50)
00:00:00
00:04:00
00:08:00
00:20:00
00:36:00
Sti-ΔCNH-GFP
D Sticky-L1246N-ΔCNH-GFP; mCh-Tubulin; sti dsRNA3 (N=56/63)
00:00:00
00:04:00
00:24:00
01:04:00
01:28:00
Sti-L1246N-ΔCNH -GFP
E Sticky-L1246N-ΔCNH-GFP; mCh-Tubulin; pbl dsRNA (N=49/55)
00:00:00
00:04:00
00:08:00
00:12:00
00:16:00
Sti-L1246N-ΔCNH -GFP
F
ns
ns
ns
Cytokinesis outcome (%)
100
80
60
40
20
0
N=149
N=57
N=52
N=60
Failure
Success
GFP-tagged construct
none
Sticky
Sti-ΔCNH
Sti-L1246N-ΔCNH
mCh-Tubulin; sti dsRNA3

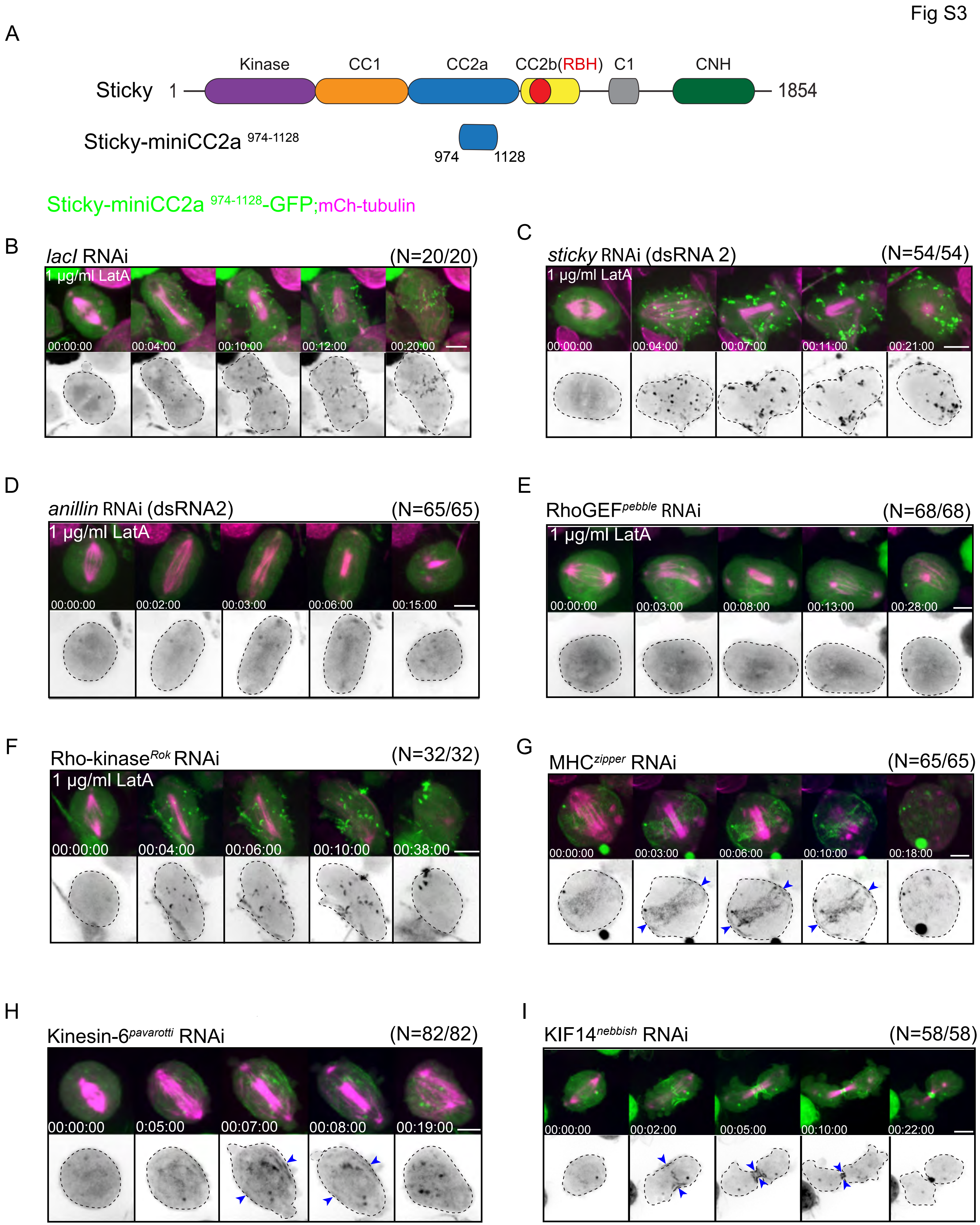
Fig S3
A
Kinase
CC1
CC2a
CC2b(RBH)
C1
CNH
Sticky 1
1854
Sticky-miniCC2a 974-1128
974
1128
Sticky-miniCC2a 974-1128-GFP;mCh-tubulin
B
lacI RNAi
(N=20/20)
1 µg/ml LatA
00:00:00
00:04:00
00:10:00
00:12:00
00:20:00
C
sticky RNAi (dsRNA 2)
(N=54/54)
1 µg/ml LatA
00:00:00
00:04:00
00:07:00
00:11:00
00:21:00
D
anillin RNAi (dsRNA2)
(N=65/65)
1 µg/ml LatA
00:00:00
00:02:00
00:03:00
00:06:00
00:15:00
E
RhoGEFpebble RNAi
(N=68/68)
1 µg/ml LatA
00:00:00
00:03:00
00:08:00
00:13:00
00:28:00
F
Rho-kinaseRok RNAi
(N=32/32)
1 µg/ml LatA
00:00:00
00:04:00
00:06:00
00:10:00
00:38:00
G
MHCzipper RNAi
(N=65/65)
00:00:00
00:03:00
00:06:00
00:10:00
00:18:00
H
Kinesin-6pavarotti RNAi
(N=82/82)
00:00:00
0:05:00
00:07:00
00:08:00
00:19:00
I
KIF14nebbish RNAi
(N=58/58)
00:00:00
00:02:00
00:05:00
00:10:00
00:22:00

Fig. S4

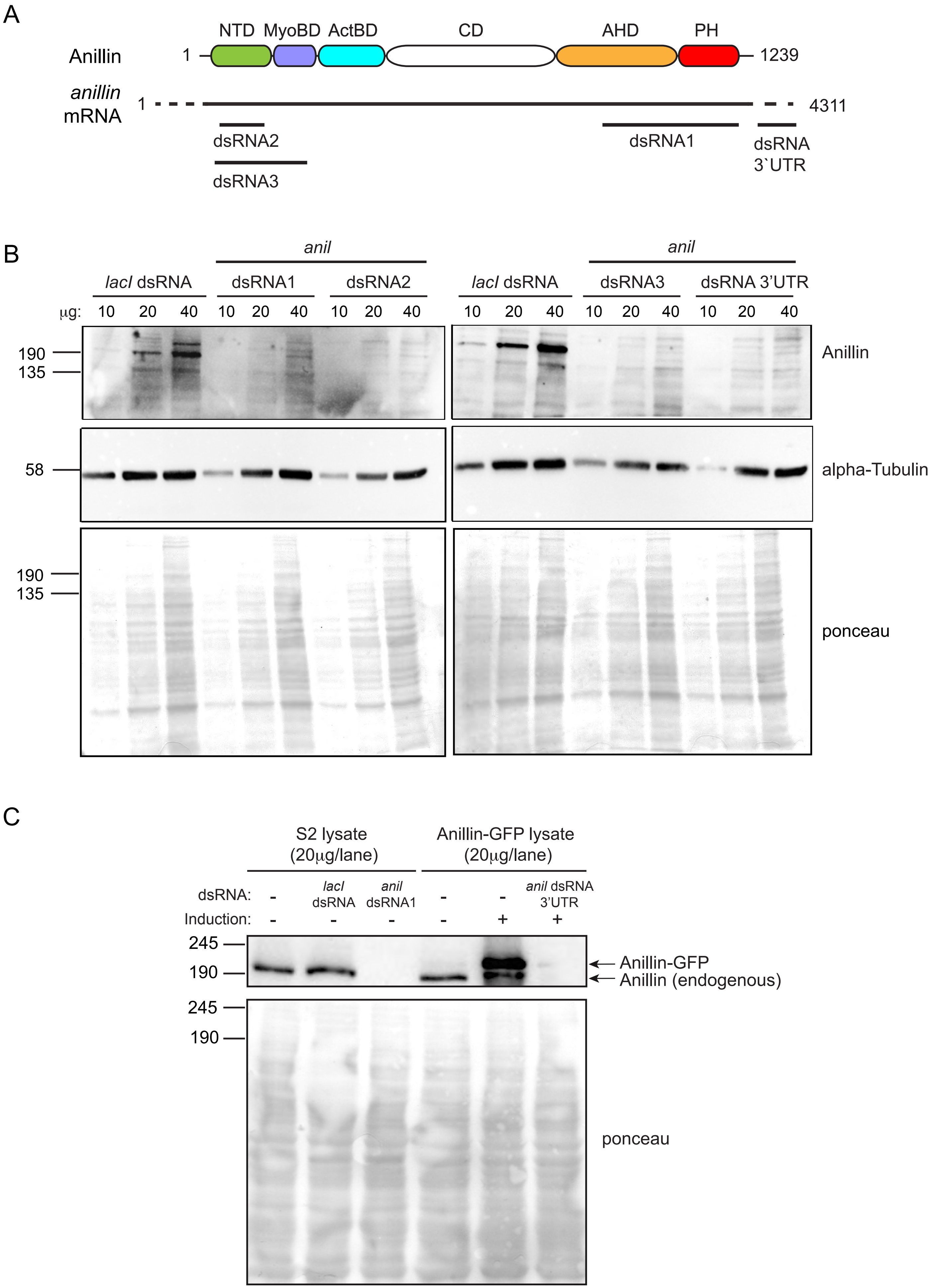
A
NTD
MyoBD
ActBD
CD
AHD
PH
Anillin
1
1239
anillin
mRNA
1
4311
dsRNA2
dsRNA1
dsRNA
3`UTR
dsRNA3
B
anil
lacI dsRNA
dsRNA1
dsRNA2
anil
lacI dsRNA
dsRNA3
dsRNA 3'UTR
μg: 10 20 40 10 20 40 10 20 40 10 20 40 10 20 40 10 20 40
190
135
Anillin
58
alpha-Tubulin
190
135
ponceau
C
S2 lysate (20μg/lane)
Anillin-GFP lysate (20μg/lane)
dsRNA: - lacI dsRNA anil dsRNA1 - - anil dsRNA 3'UTR
Induction: - - - - + +
245
190
Anillin-GFP
Anillin (endogenous)
245
190
ponceau

Fig. S5

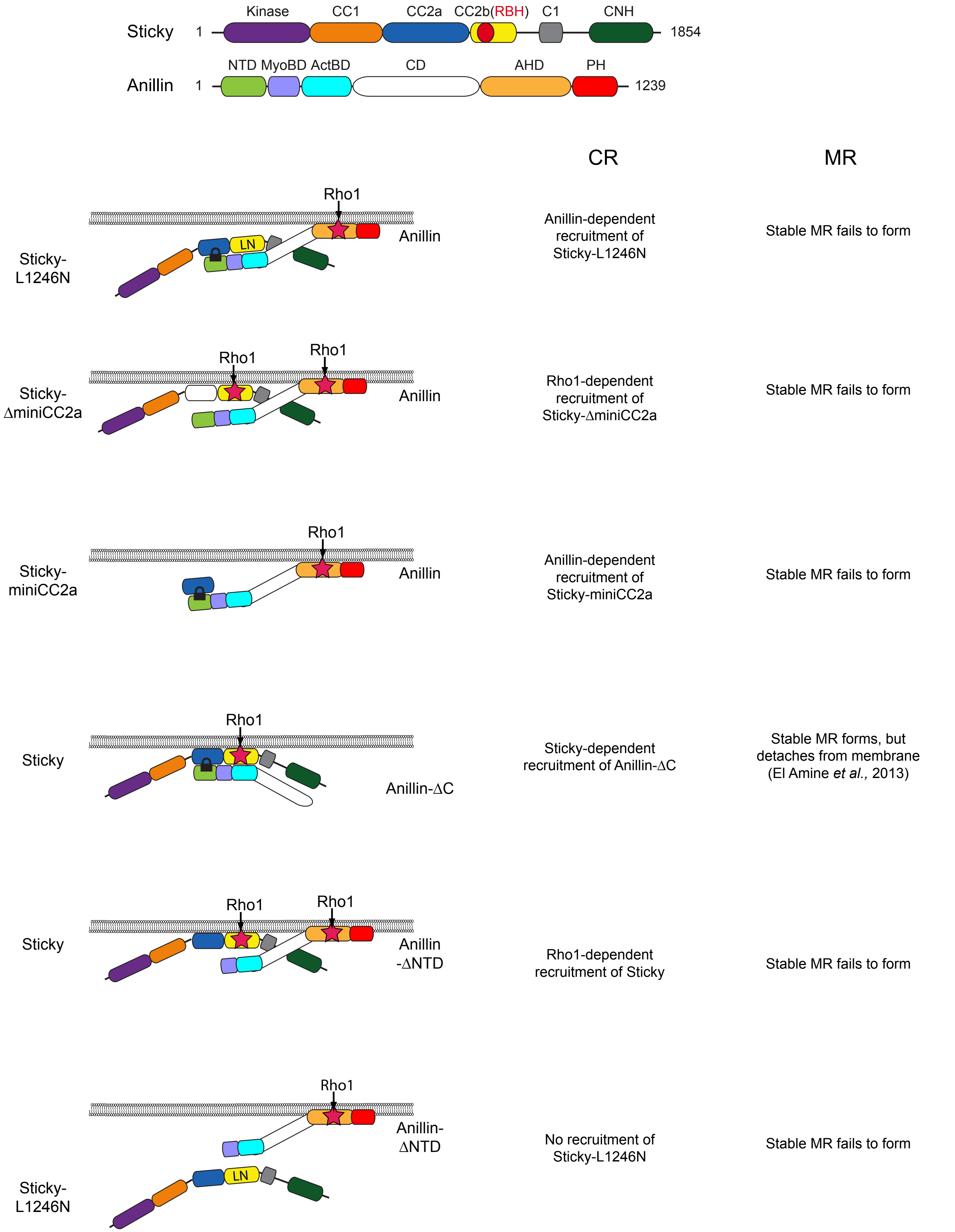

Sticky
1
Kinase
CC1
CC2a
CC2b(RBH)
C1
CNH
1854
Anillin
1
NTD MyoBD ActBD
CD
AHD
PH
1239
CR
MR
Sticky-L1246N
Rho1
LN
Anillin
Anillin-dependent recruitment of Sticky-L1246N
Stable MR fails to form
Sticky-ΔminiCC2a
Rho1
Rho1
Anillin
Rho1-dependent recruitment of Sticky-ΔminiCC2a
Stable MR fails to form
Sticky-miniCC2a
Rho1
Anillin
Anillin-dependent recruitment of Sticky-miniCC2a
Stable MR fails to form
Sticky
Rho1
Anillin-ΔC
Sticky-dependent recruitment of Anillin-ΔC
Stable MR forms, but detaches from membrane (El Amine *et al.*, 2013)
Sticky
Rho1
Rho1
Anillin -ΔNTD
Rho1-dependent recruitment of Sticky
Stable MR fails to form
Rho1
Anillin-ΔNTD
LN
Sticky-L1246N
No recruitment of Sticky-L1246N
Stable MR fails to form
